# Leadership of supported and new clinical trials after the NIH K23 clinical trial pathway

**DOI:** 10.64898/2026.10.09.758109

**Authors:** Haining Wang, Hengyi Fu, Zuoyu Tian

## Abstract

**Background:** In 2018, the US National Institutes of Health (NIH) created a dedicated clinical trial pathway for K23 career development awards, and its planned KC1 award will not fund trial conduct for applications due from late 2027. Whether the pathway changed how many awardees led trials, or which trials, is unknown. We assessed leadership of the trial the award supported (K trial), of distinct new trials, and of any trial.

**Methods and findings:** In this cohort study of 1847 K23 awardees (fiscal years 2014 to 2021; 837 under the pathway), we linked NIH award records to ClinicalTrials.gov registrations, including archived versions. A difference-in-differences design, adjusted for fiscal year and NIH funding institute, compared awardees whose funded abstracts proposed their own trial (trial plans; n = 721) with other awardees. A language model measured trial plans and leadership, and author review of 280 sampled records corrected its errors. Before correction, 54.4% of earlier and 73.1% of pathway awardees with trial plans led a K trial within 5 years, vs 4.1% and 5.1% of others. This gap widened by 18.6 percentage points uncorrected and 13.7 (95% CI, 5.7-21.2) corrected, mostly in grant-citing trials. The new-trial gap changed by *−*3.8 (95% CI, *−*14.4 to 6.9). Assuming parallel trends, every correction excluded gains above 11.5 points (earlier rate, 37.6%). The any-trial gap widened by 4.2 (95% CI, *−*5.9 to 14.0) when correction probabilities were estimated separately by period and 9.6 (95% CI, 1.5-17.9) when they were pooled across periods. Had early-cohort trends (2.4 points yearly) continued, the K-trial estimate would be 4.1 (95% CI, *−*17.7 to 26.8). Limitations include more complete registration after concurrent NIH application changes, which the design cannot separate from the pathway, and selection among funded awardees.

**Conclusions:** The dedicated pathway was associated with more awardees with trial plans leading their award-supported trial, although more complete registration or an earlier trend could not be ruled out. New-trial leadership within 5 years did not detectably increase, and whether the pathway produced more trial leaders is unclear. Evaluations of KC1 should track proposed and new trials separately.

**Author summary:** *Why was this study done?:*

- The US National Institutes of Health (NIH) trains clinician investigators through K23 career development awards. In 2018, NIH split these awards into a pathway that required applicants to propose leading their own clinical trial and one that did not allow it.
- NIH will consolidate its clinician K awards into a single KC1 award for applications due from late 2027, and KC1 funds cannot pay for trials, yet little is known about what the dedicated pathway did.

*What did the researchers do and find?:*

- We followed 1847 K23 awardees for 5 years in ClinicalTrials.gov, comparing awardees whose funded plans proposed their own trial with other awardees before and after the pathway, and corrected language-model measurements with author review of 280 records.
- In registry records, more awardees with trial plans led the trial their award supported after the pathway (73.1% vs 54.4% before correction); after correction, the gap with other awardees widened by 13.7 percentage points (95% CI, 5.7-21.2).
- Leadership of distinct new trials did not detectably increase, and whether more awardees led any trial depended on an assumption about measurement error.
- The increase came mostly from registrations citing the K grant, and more complete registration after concurrent NIH changes or a continuing earlier trend could not be ruled out.

*What do these findings mean?:*

- A dedicated trial pathway coincided with more awardees leading their award-supported trial, but not with detectably more awardees leading new trials within 5 years of award start.
- In uncorrected registry records, 69.9% of the 346 pathway awardees with trial plans had started a K trial and 39.6% had one recorded as completed within 5 years, a baseline for evaluating KC1.
- Because KC1 awards cannot fund trials, registry grant citations may no longer identify awardees’ trials; evaluations should record each trial’s funder and whether the award plan proposed it, and follow awardees for 7 years.

## Introduction

Clinical trials need clinicians who can design and lead them, yet physicians now make up a smaller share of first-time National Institutes of Health (NIH) R01-equivalent recipients than in the 1980s [1]. NIH trains clinician investigators through mentored career development (K) awards, and from late 2027, K award funds will no longer pay for clinical trials [2]. Funders support projects partly to develop investigators who will lead further work, and delivering the funded project and leading new work are distinct outcomes [3, 4]. Studies of NIH funding have also shown that changes in who is funded can look like program effects [5–7].

K23 awards for patient-oriented research offer a setting in which these outcomes can be separated. Earlier K23 awards allowed but did not require awardees to lead their own trials. For applications due from January 25, 2018, NIH split K23 awards into a pathway that required applicants to propose leading a trial within the award, with trial-specific review, and a pathway that did not allow one (Box 1). NIH framed the change within its stewardship of clinical trials rather than as a career development aim [8].

### Box 1. The K23 clinical trial pathway and KC1

- **Earlier announcements (until January 2018).** K23 awards support mentored patient-oriented research for a project period that may not exceed 5 years. Awardees could lead their own trial within the award, but none was required [9, 10].
- **Independent Clinical Trial Required (PA-18-374, PA-19-118, and PA-20-206; from January 25, 2018).** Applicants proposed a trial they would lead within the award, and review applied trial-specific criteria [11]. Institutes listed $25,000 to $51,000 per year for research costs [12].
- **Independent Clinical Trial Not Allowed (PA-18-375, PA-19-119, and PA-20-205; from January 25, 2018).** Awardees could not lead their own trial with the award. They could gain experience in a mentor’s trial, and their own trials required separate support [11].
- **KC1 (applications due from October 12, 2027).** One award replaces K08, K23, and K38, and awardees who wish to lead a trial “must secure separate funding” [2].

Leading a distinct new trial is one visible outcome that could follow mentored trial experience, although it does not directly measure trial skills or independence [13]. If the pathway developed trialists, more awardees with trial plans could have led new trials within 5 years of award start. If it instead paid for trials that awardees would have led anyway with other funds, or favored applicants whose trials were ready to start, more awardees should have led the award-supported trial (K trial), but not more new trials. We therefore asked whether K23 awardees with trial plans became more likely to lead award-supported trials, new trials, or any trial after the dedicated pathway.

## Methods

This cohort study used only public records of NIH awards and registered trials. It is reported following the Strengthening the Reporting of Observational Studies in Epidemiology (STROBE) guideline (S1 Checklist).

### Data sources and cohorts

We queried NIH RePORTER for type 1 K08 and K23 awards. The earlier cohort comprised 1860 awards starting from fiscal year 2014 under parent announcements PA-11-193, PA-11-194, PA-14-046, PA-14-049, PA-16-191, and PA-16-198. The pathway cohort comprised 1527 awards starting from January 1, 2018, through September 27, 2021, under the Required and Not Allowed parent announcements (S1 Appendix). Each investigator contributed the earliest qualifying award, no investigator was in both cohorts, and all awardees had at least 5 years of follow-up by September 2026. An abstract classifier excluded 25 basic-experimental-only projects, leaving 3362 awardees, of whom 1847 held K23 awards (Fig 1). K23 awardees were the study population. K08 estimates are reported separately because only 14 earlier K08 awardees planned trials.

**Fig 1.**
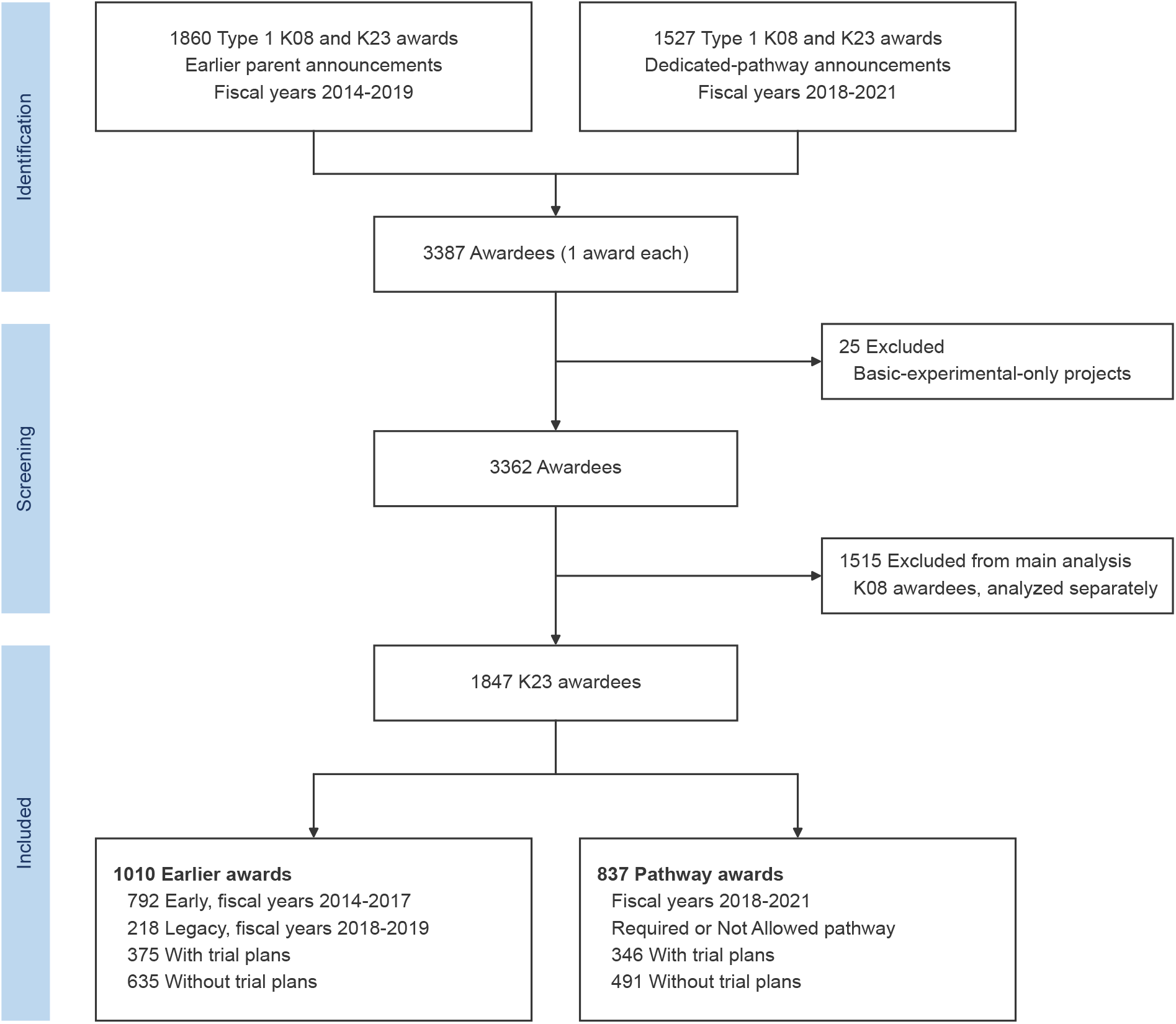
Study population. Earlier awards started in fiscal years 2014 to 2019 under the earlier parent announcements; legacy awards were those starting in fiscal years 2018 to 2019. Each investigator contributed the earliest qualifying award, and no investigator was in both cohorts. Trial plans were classified from the funded abstract. NIH indicates National Institutes of Health.

### Exposure, comparison groups, and estimand

The exposure was the announcement type. Early awards started in fiscal years 2014 to 2017. Legacy awards started in fiscal years 2018 to 2019 (nearly all in 2018) under the earlier announcements, which expired on January 25, 2018. Trial intent, hereafter trial plans, was classified from the funded abstract, with funding-designation phrases removed, as a plan for a prospective interventional study led by the awardee. The estimand is the change, from earlier to pathway awards, in the leadership gap between funded awardees with and without trial plans. Because the pathway could change who received awards and how plans were written, this estimand combines effects on awardees with changes in who was funded. Most pathway awardees without trial plans held Not Allowed awards, so the comparison group was also exposed to the policy change.

### Outcomes

Candidate trials were retrieved from ClinicalTrials.gov by investigator name, grant number, and surname with institution, including archived records from 2022 and 2024 [14, 15]. Leadership required the awardee to be listed as overall principal investigator (PI), or as responsible party of investigator type, for an interventional trial testing a clinical or care intervention that started, by actual or anticipated date, within 5 years of K start, without requiring evidence of enrollment.

Because a K23 project period may not exceed 5 years [9], this window falls mostly within the award. Registry-documented K-trial leadership required a trial that listed the K grant in the registry identification fields, which classified it as a K trial regardless of content, or that matched the proposed K study. This category is operational: a grant citation or plan match does not show how much the K award paid for the trial. Other qualifying trials were new trials, and any-trial leadership meant leading either type. Each awardee was also classified by the combination of trials led: neither trial type, a K trial only, a new trial only, or both. Secondary outcomes were registration with the awardee as PI and a K-grant link, recorded initiation, results submission within 12 months of primary completion, new-trial leadership by 7 years, new trials starting 3 or more years after K start, and receipt of an R01-equivalent award [16].

### Measurement and human validation

A locally run language model (Qwen/Qwen3.8-27B-FP8) classified trial plans and assessed the identity, role, scope, and relation to the K plan of each candidate trial. Two authors (H.F. and Z.T.) reviewed a random sample of 280 K23 awardee records, stratified by award period, trial plans, and model-assigned state, using ClinicalTrials.gov and their own name searches (Table A in S1 Appendix). They did not see announcement labels or model output, but they saw award dates, so timing was inferable. Both rated 110 records independently, and one rated each of the other 170. Reference labels came from H.W.’s resolution of disagreements for 70 double-rated records and from a preassigned rater for 40. Further samples covered trial plans in 100 abstracts and the K-trial and new-trial boundary in 60 pairs (Table B in S1 Appendix). A second model (GPT-5.6 Sol) rated plan continuity (S1 Appendix). The trial combinations of unrated awardees were imputed from Dirichlet distributions within cells defined by award period (early, legacy, or pathway), trial plans, and model-assigned state, with a centered prior placing 0.5 on the model state and 0.01 on each alternative and with validation counts fixed across bootstrap draws [17]. Within each cell, rated and unrated awardees were assumed exchangeable, and the distribution of human labels given the model state (the correction probabilities) was assumed not to vary with fiscal year or institute. Cells without rated awardees borrowed counts from a coarser parent cell (S1 Appendix). Because error that differs by period would bias a difference-in-differences (DID) estimate, and among awardees with trial plans the model called a K trial for 4 of 18 pathway and 0 of 18 early records the reviewers judged negative (unweighted counts; Table B in S1 Appendix), the primary correction estimated the correction probabilities separately by period.

### Statistical analysis

Linear probability models regressed each outcome on trial plans, announcement type, and their interaction, with fiscal-year and NIH funding institute fixed effects. The interaction is the adjusted DID estimate. The primary estimates were human-corrected DIDs for K-trial, new-trial, and any-trial leadership, computed as means over 2000 bootstrap draws that resampled the 185 institutions and repeated the correction, with percentile 95% CIs. The bootstrap accounts for institutional clustering of outcomes but not of classification errors. DIDs for the trial combinations are linear combinations of these 3. For example, the new-trial-only DID equals the any-trial minus the K-trial DID. Within the same draws, we fitted the models with legacy awards as the comparator (153 institutions) and with both announcement types restricted to fiscal years 2018 to 2019. A trend check subtracted from the outcomes of awardees with trial plans, in every fiscal year, the early-cohort difference in yearly slope between awardees with and without trial plans, estimated in fiscal year 2014 to 2017 cohorts with the same fixed effects and multiplied by fiscal year minus 2017. We also report the annual drift in the gap after fiscal year 2017 at which the K-trial interval would reach zero. A drift of *d* percentage points per year moves each interval limit outward by *d* multiplied by the DID that an outcome equal to trial plans multiplied by years after fiscal year 2017 would yield (2.76 years on average). This check addresses gradual change, not a step at the policy date. Sensitivity analyses varied the correction prior, rater, correction of trial plans, and pooling of correction probabilities across periods (7 alternatives to the primary correction). Composition analyses (covariate adjustment, inverse probability weighting, and trimming on prior trials, prior NIH awards, and institution size), an analysis excluding trial starts in 2020 to 2021, which removes more follow-up from pathway awards and so is biased against them (Table F in S1 Appendix), and secondary outcomes used uncorrected labels and 1000 draws. Intervals are pointwise, and secondary analyses were not adjusted for multiple comparisons. Analyses used Python, version 3.12 (Python Software Foundation), and R, version 4.6 (R Foundation for Statistical Computing).

### Ethics statement

Because the study did not involve interaction with individuals or identifiable private information, it did not meet the definition of human participants research under 45 CFR 46.102, and institutional review board review and informed consent were not required. Results are reported in aggregate, and no individual investigator is identified.

### Use of artificial intelligence

Qwen/Qwen3.8-27B-FP8 was the locally served measurement instrument, and GPT-5.6 Sol provided model lineage ratings assessed against human references. In addition, the authors used Claude Fable 5.1 solely for copyediting, reviewed and revised the text as needed, and take full responsibility for its content.

### Data and code availability

All source data are public NIH RePORTER and ClinicalTrials.gov records. Code for every analysis, model prompts, the rating codebook, aggregate results, and pseudonymized awardee-level analysis files, which include the human reference labels, are available at https://github.com/ Wang-Haining/k under the MIT license. These files reproduce the uncorrected leadership estimates. Confidence intervals, the corrected estimates (means over institution-bootstrap draws), and the analyses of prior NIH awards, institution size, and trial characteristics also need institution identifiers, award histories, or the trial-level linkage. These files and the completed rating workbooks are not posted because they identify investigators. The repository code can retrieve the public records again, which yields a new snapshot rather than the frozen inputs used here.

## Results

### Study population

Of 1847 K23 awardees, 1010 held earlier awards (792 early and 218 legacy) and 837 held pathway awards. In total, 721 had trial plans (Table 1). Demographic characteristics are not available in public award records. Prior non-K trial leadership was similar across announcements (18.1% vs 20.2% among awardees with trial plans), but fewer pathway awardees without trial plans held prior NIH awards as PI (16.5% vs 21.6%; DID, 7.5 percentage points; 95% CI, 0.1-15.2). Among contemporaneous K23 awardees, 358 of 825 held Required awards, and trial plans appeared in 90.2% of Required and 5.1% of Not Allowed abstracts (Table C in S1 Appendix).

**Table 1.**
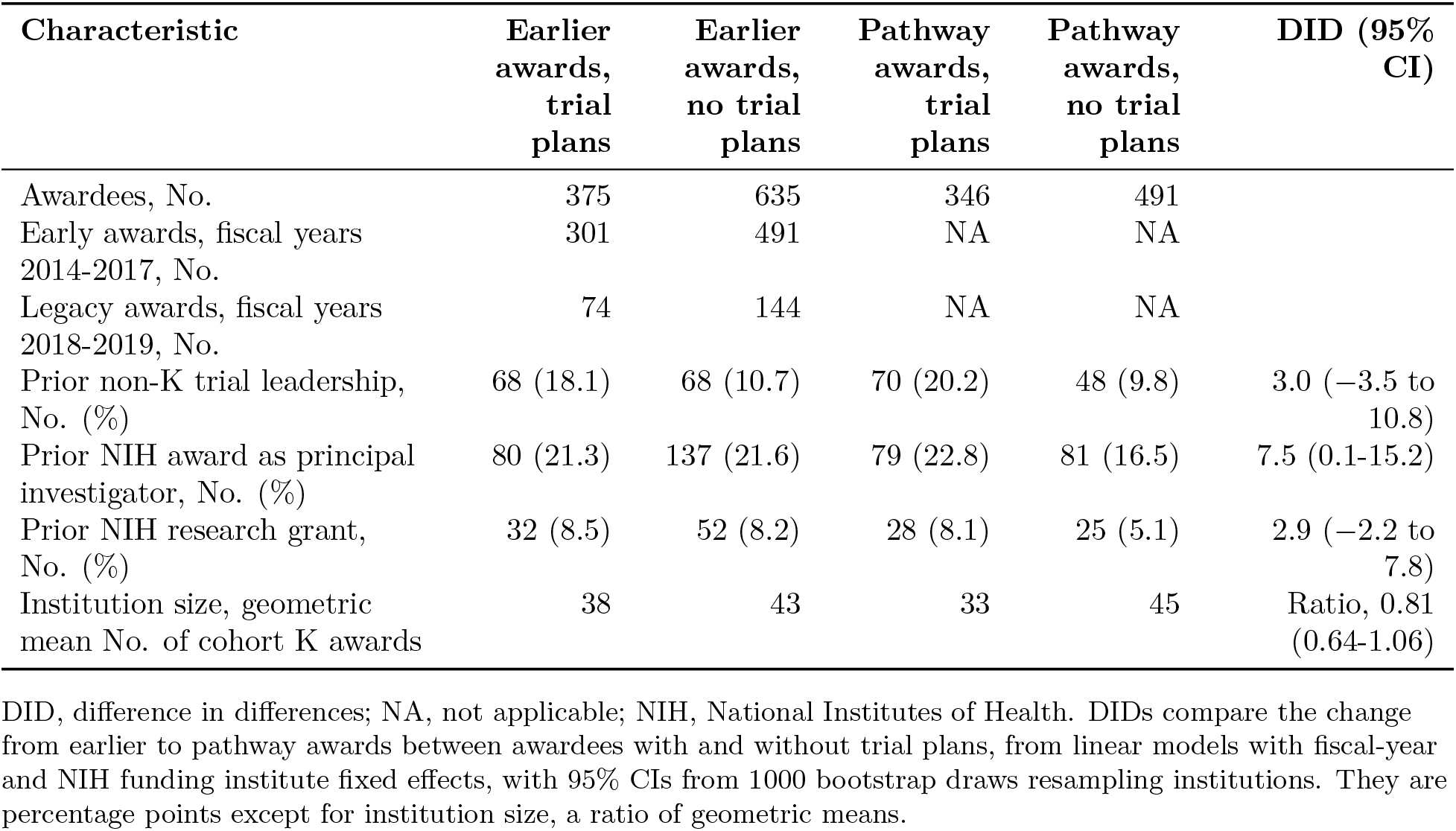
Characteristics of K23 awardees by announcement and trial plans.

| Characteristic | Earlier<br>awards,<br>trial<br>plans | Earlier<br>awards,<br>no trial<br>plans | Pathway<br>awards,<br>trial<br>plans | Pathway<br>awards,<br>no trial<br>plans | DID (95%<br>CI) |
| --- | --- | --- | --- | --- | --- |
| Awardees, No. | 375 | 635 | 346 | 491 |  |
| Early awards, fiscal years<br>2014-2017, No. | 301 | 491 | NA | NA |  |
| Legacy awards, fiscal years<br>2018-2019, No. | 74 | 144 | NA | NA |  |
| Prior non-K trial leadership,<br>No. (%) | 68 (18.1) | 68 (10.7) | 70 (20.2) | 48 (9.8) | 3.0 (−3.5 to<br>10.8) |
| Prior NIH award as principal<br>investigator, No. (%) | 80 (21.3) | 137 (21.6) | 79 (22.8) | 81 (16.5) | 7.5 (0.1-15.2) |
| Prior NIH research grant,<br>No. (%) | 32 (8.5) | 52 (8.2) | 28 (8.1) | 25 (5.1) | 2.9 (−2.2 to<br>7.8) |
| Institution size, geometric<br>mean No. of cohort K awards | 38 | 43 | 33 | 45 | Ratio, 0.81<br>(0.64-1.06) |

DID, difference in differences; NA, not applicable; NIH, National Institutes of Health. DID's compare the change from earlier to pathway awards between awardees with and without trial plans, from linear models with fiscal-year and NIH funding institute fixed effects, with 95% CIs from 1000 bootstrap draws resampling institutions. They are percentage points except for institution size, a ratio of geometric means.

### Trial leadership

Before correction, 54.4% (204/375) of earlier and 73.1% (253/346) of pathway awardees with trial plans led a K trial, compared with 4.1% (26/635) and 5.1% (25/491) of awardees without trial plans (Fig 2). Because few awardees without trial plans led a K trial, the K-trial DID mainly reflects change among awardees with trial plans. After correction, the gap widened by 13.7 percentage points (95% CI, 5.7-21.2) for K-trial leadership and changed by *−*3.8 (95% CI, *−*14.4 to 6.9) for new-trial leadership. Across all 8 corrections, the K-trial DID ranged from 10.3 to 16.8 (lower limits, *−*1.7 to 9.7; 7 of 8 excluded zero) and the upper limit for new-trial gains from 5.5 to 11.5 percentage points, against an earlier new-trial rate of 37.6% (Table D in S1 Appendix). Any-trial leadership changed by 4.2 (95% CI, *−*5.9 to 14.0) when correction probabilities were estimated separately by period, 9.6 (95% CI, 1.5-17.9) when they were pooled across periods, and 10.8 (95% CI, 2.2-19.2) before correction. All 8 corrections gave positive estimates (1.5 to 9.6), but 7 of 8 intervals included zero. Most of the decline in awardees leading only a new trial (*−*9.4; 95% CI, *−*16.7 to *−*2.1) was offset by more awardees leading both trial types (5.7; 95% CI, *−*1.0 to 12.5). Adjusting or reweighting for prior trials, prior NIH awards, and institution size barely changed the uncorrected K-trial and new-trial DIDs of 18.6 and *−*2.8 (Table E in S1 Appendix).

**Fig 2.**
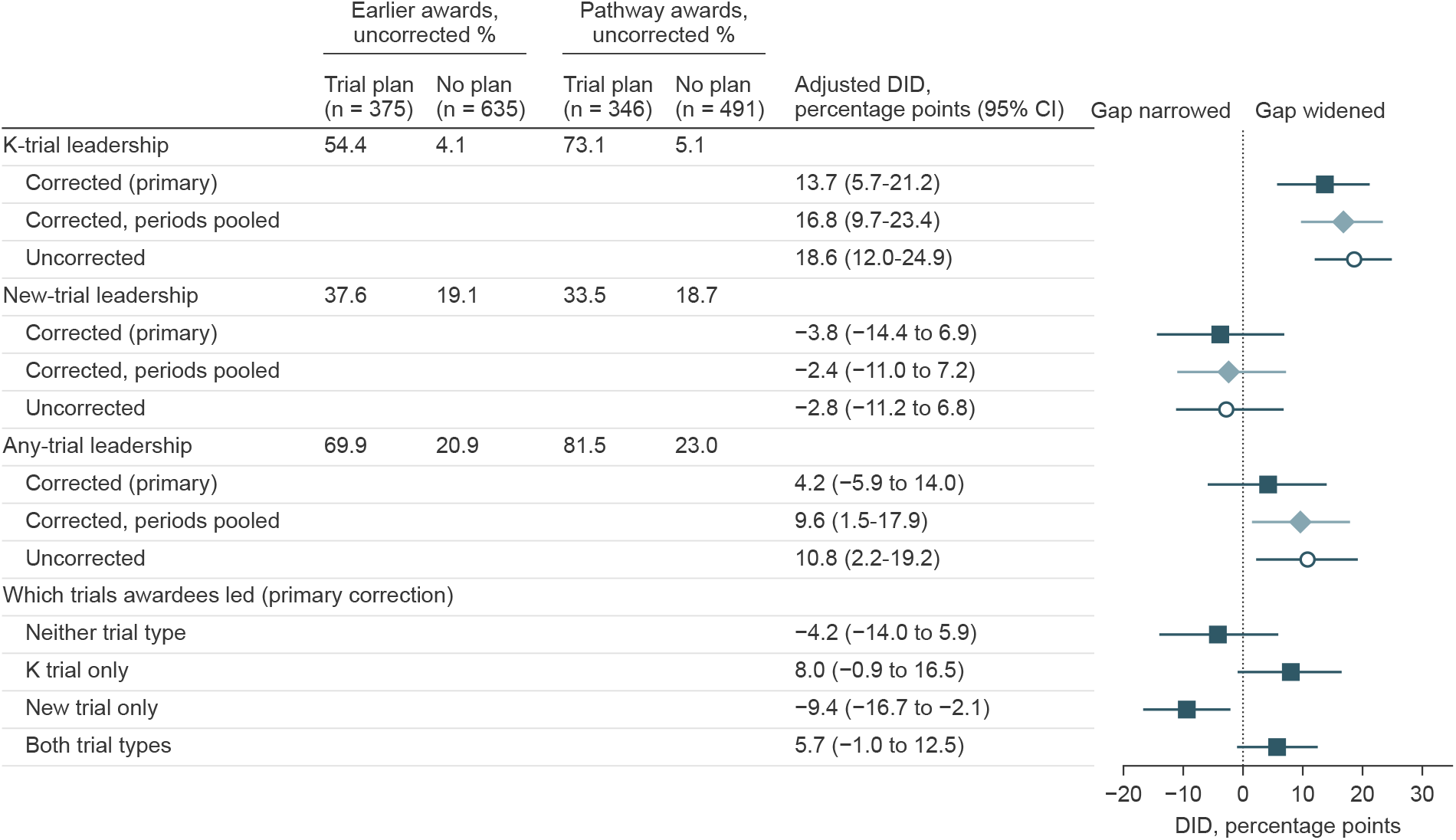
Leadership of award-supported (K) trials, new trials, and any trial among K23 awardees. Percentages are uncorrected registry-documented leadership within 5 years of K award start among awardees with and without trial plans. Difference-in-differences (DID) estimates, adjusted for fiscal year and NIH funding institute, show how much the gap between these groups changed from earlier to pathway awards; positive values mean the gap widened. Squares indicate the primary correction, which estimates correction probabilities separately by period; diamonds, the correction pooling them across periods; and circles, uncorrected estimates. Corrected estimates are means of 2000 bootstrap draws resampling institutions, and uncorrected estimates are full-sample fits; whiskers show 95% CIs. The 4 combinations of trials led are shown for the primary correction only; their DIDs are linear combinations of the K-trial, new-trial, and any-trial DIDs and sum to zero before rounding. Part of the decline in leading only a new trial is arithmetic: if awardees who newly led a K trial had led new trials at the uncorrected earlier rate (37.6%), about 5 points would move from new trial only to both without any displacement. The new-trial-only interval included zero under 2 of the 8 corrections (Table D in S1 Appendix).

### Comparison group and trend checks

Comparisons built on legacy awards gave nearly the same K-trial DID with wider intervals: 13.5 percentage points (95% CI, *−*0.3 to 27.3) against all pathway awards and 12.9 (95% CI, *−*3.9 to 29.1) within fiscal years 2018 to 2019 (Fig 3; Tables F and G in S1 Appendix). Among awardees with trial plans, 47.2% to 59.5% of each early cohort and 54.2% of 2018 legacy awardees led a K trial, vs 70.7% to 77.1% of each pathway cohort from 2019 (Fig 4). Extrapolating early-cohort trends gave a K-trial DID of 4.1 (95% CI, *−*17.7 to 26.8), too imprecise to confirm or exclude the increase. Had the gap widened by about 2 percentage points per year after 2017 for other reasons, the K-trial interval would include zero. In early cohorts it widened 2.4 points per year (95% CI, *−*3.0 to 7.9), and an uncorrected early-cohort placebo gave 7.7 (95% CI, *−*1.7 to 19.2). Neither excludes such a trend.

**Fig 3.**
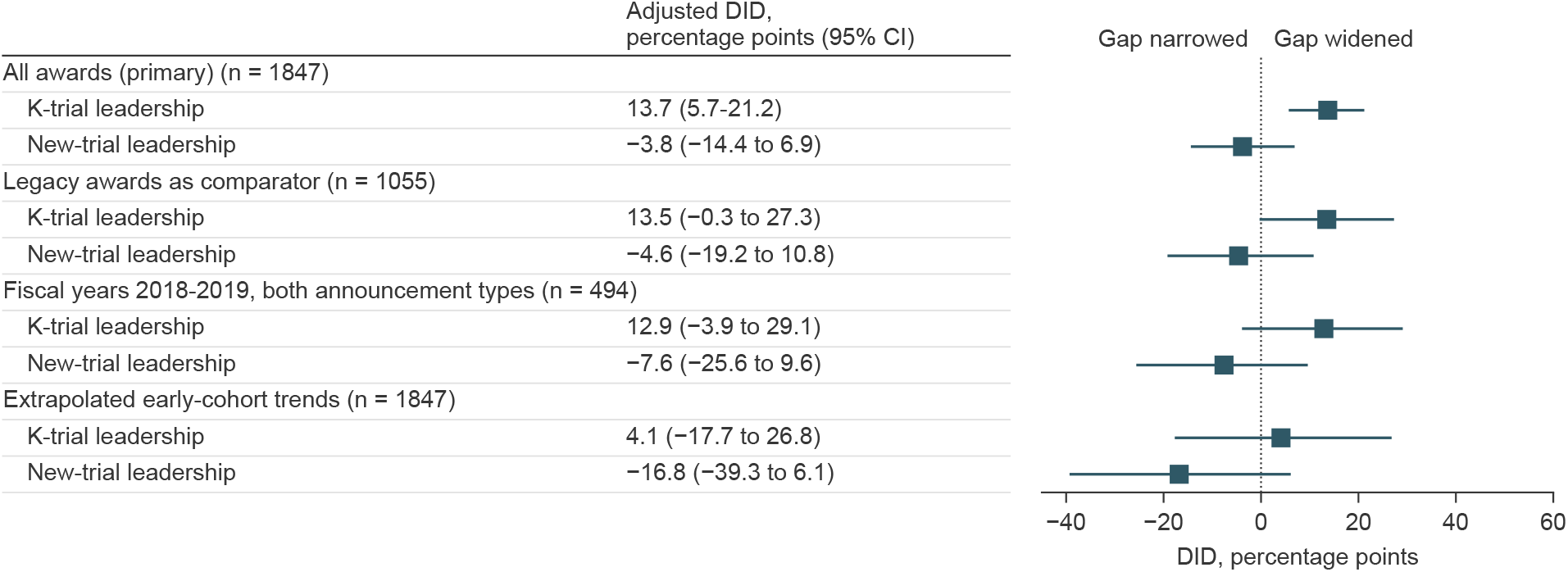
Comparison-group and trend checks. Difference-in-differences (DID) estimates, corrected and adjusted as in the primary analysis, from the same 2000 bootstrap draws. The legacy comparison replaces all earlier awards with legacy awards; the fiscal year 2018 to 2019 comparison restricts both announcement types to those years and rests mainly on 72 legacy awardees from 2018 and 89 pathway awardees from 2019 with trial plans; the trend row subtracts linear trends by trial plans estimated in fiscal year 2014 to 2017 cohorts. Whiskers show 95% CIs.

**Fig 4.**
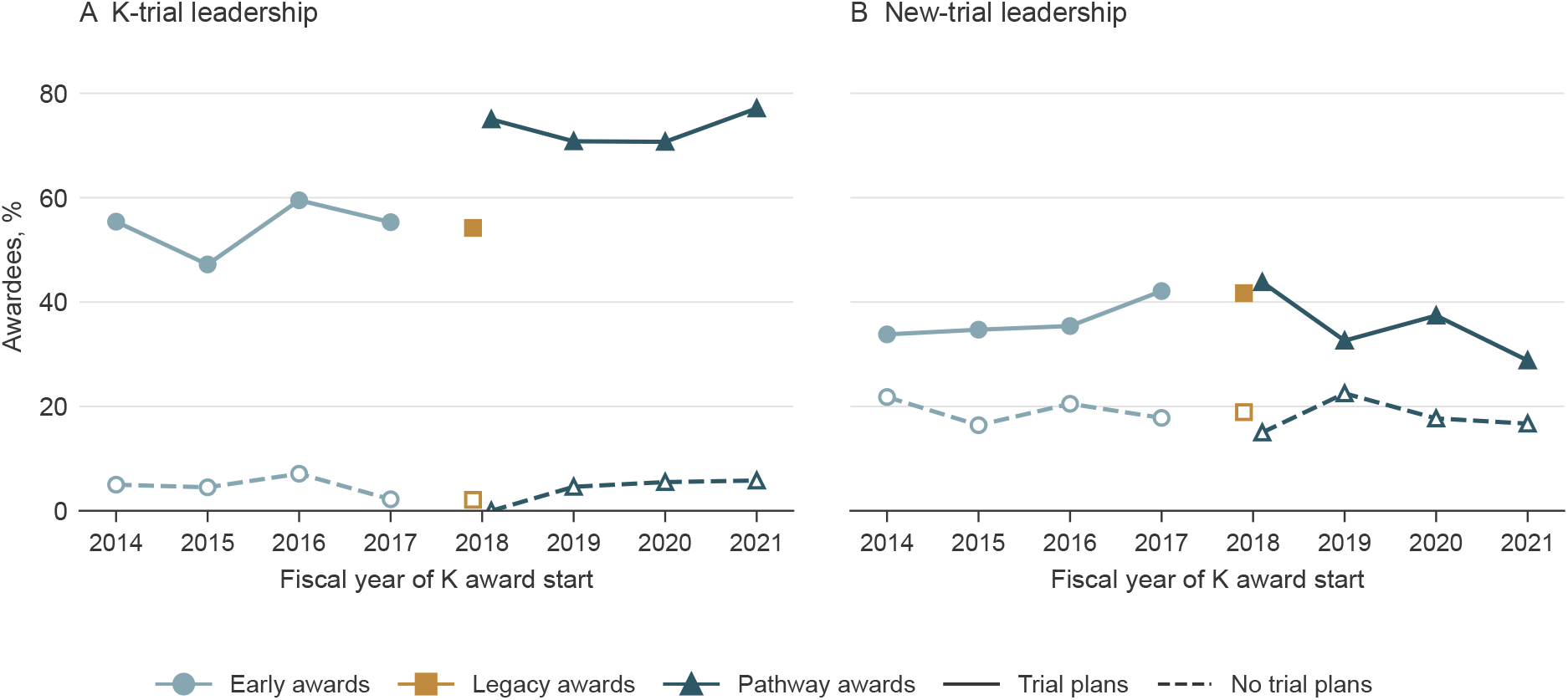
Annual leadership of award-supported (K) trials and new trials among K23 awardees. Registry-documented leadership within 5 years of K award start, as measured by the language model, by fiscal year of award start. Solid lines indicate awardees with trial plans and dashed lines, awardees without trial plans. Fiscal year 2018 pathway points include 16 awardees with and 20 without trial plans, and fiscal year 2019 legacy cells with fewer than 10 awardees are omitted. Annual rates are descriptive and uncorrected. Each point is an award cohort; the 5-year follow-up of earlier cohorts extends into the pathway years, so earlier points are not outcomes measured before the policy.

### Registration, trial characteristics, and reporting

The increase came from trials whose registry record cited the K grant (uncorrected DID, 17.8; 95% CI, 10.8-24.8), not from trials matched only to the K plan (0.8; 95% CI, *−*4.2 to 6.0). Grant-citing trials were led by 40.5% of early, 44.6% of legacy, and 59.2% of pathway awardees with trial plans, while plan-matched-only shares stayed at 14.9% to 17.3% (Table H in S1 Appendix). New-trial leadership defined by scientific distinctness alone, ignoring grant links, also did not detectably increase (*−*3.1; 95% CI, *−*11.9 to 6.3) (Table E in S1 Appendix). Among awardees with trial plans, later addition of a K link increased relative to early awards (7.9; 95% CI, 2.7-13.3) and did not differ detectably from legacy awards (*−*1.0; 95% CI, *−*11.1 to 7.0) (Fig A in S1 Appendix). Evidence that a K trial had started (recorded recruitment or actual enrollment) was documented for 52.8% of early, 52.7% of legacy, and 69.9% of pathway awardees with trial plans, and a K trial recorded as completed, with primary completion within 5 years, for 30.9%, 29.7%, and 39.6%. K trials were small and rarely run at 3 or more sites in both periods (median enrollment, 60 and 58; 3 or more sites, 3.9% and 5.5%). Pathway K trials were less often randomized (68.0% vs 77.9%) and, with less time since starting, more often ongoing (22.5% vs 1.5%) (Table I in S1 Appendix). Of K trials due to report, 25.5% of early, 45.0% of legacy, and 46.5% of pathway trials submitted results within 12 months of primary completion (Fig B and Table J in S1 Appendix).

### Outcomes beyond the supported trial

Among 1274 awardees followed 7 years (264 with pathway awards), counting only trials registered by each time point, the uncorrected new-trial DID was *−*6.3 percentage points (95% CI, *−*18.2 to 5.2) at 5 years and *−*3.4 (95% CI, *−*16.7 to 8.9) at 7 years (Table K in S1 Appendix). New trials starting 3 or more years after K start (*−*1.4; 95% CI, *−*8.4 to 6.8) and R01-equivalent awards within 5 years, an early transition (*−*0.7; 95% CI, *−*8.1 to 6.7) showed no detectable increase, and K08 estimates were imprecise (Table L in S1 Appendix).

## Discussion

In this cohort study of K23 awardees, NIH’s dedicated clinical trial pathway was associated with more registry-documented leadership of award-supported trials and no detectable increase in new-trial leadership within 5 years. The K-trial increase was positive under every correction, with 7 of 8 intervals excluding zero, and comparisons built on legacy awards agreed in size, but more complete registration or a continuing earlier trend could not be ruled out. The any-trial answer depended on whether the relation between model calls and human labels differed by period, which the validation sample could not settle, so it is unclear whether the pathway produced more trial leaders. Assuming parallel trends, new-trial gains of up to 11.5 percentage points remain possible and would matter for workforce development. This limit, like every estimate here, includes changes in who was funded. NIH framed the pathway as trial stewardship. New-trial leadership bears on the workforce aim that KC1 now raises.

Several explanations fit these results. If the pathway supported trials that awardees planned and would not otherwise have run, more awardees would lead any trial. If K funds paid for trials that awardees would otherwise have led as new trials with other funds, K-trial leadership would rise and new-trial leadership fall without more leaders. Registry records cannot distinguish these 2 explanations because they do not show which funds paid for trial conduct. If running the K trial used protected time and delayed separate trials, new trials would appear later. Among awardees followed 7 years, the uncorrected new-trial DID moved from *−*6.3 at 5 years to *−*3.4 at 7 years, too imprecise to separate delay from substitution. If awardees cited the K grant in the registry more often, some trials would count as K trials rather than new trials through documentation alone. New-trial leadership defined by content alone, which ignores grant links, did not detectably increase either (uncorrected upper confidence limit, 6.3 points). Trials matched to the K plan without a grant citation did not become rarer, so if better documentation explains the rise, it involves trials earlier awardees never registered, never linked, or registered only under a mentor’s name. Trial-specific review may also have selected applicants whose trials were ready to start, and such selection is part of what the policy did. Because this study compares funded awardees rather than applicants [5, 6], its estimates include this selection. It complements work on K-supported trials and later independence [18–20].

Two other NIH changes applied to applications due from January 25, 2018: trial-specific funding announcements that followed the revised clinical trial definition and new application forms collecting clinical trial information. Both could have increased registration and grant listing of K trials. This explanation, rather than more awardees leading trials, is the one these data can least rule out [8, 21, 22]. The Food and Drug Administration Amendments Act final rule and NIH dissemination policy likely applied to legacy and pathway awards alike, as the rise in timely reporting from legacy awards suggests, so they are unlikely to explain the legacy comparison [23, 24]. The fiscal year 2018 to 2019 comparison holds calendar time, including the COVID-19 pandemic, roughly constant and gave a similar K-trial DID (12.9; 95% CI, *−*3.9 to 29.1).

As a baseline for KC1, uncorrected registry records showed that 69.9% (242/346) of pathway awardees with trial plans had started a K trial and 39.6% (137/346) had one recorded as completed within 5 years, vs 52.8% (159/301) and 30.9% (93/301) of early awardees. Under KC1, such small trials, rarely run at 3 or more sites, will need separate funding, and whether early-career investigators can obtain it is unknown. The Not Allowed pathway already followed these rules, but its recipients chose it, so it cannot show how awardees who would have chosen the Required pathway will fare without trial funds. Because KC1 awards cannot fund trials, registry grant citations may no longer identify awardees’ own trials. Progress reports could instead list each trial an awardee leads, its funder, and whether the award plan proposed it, and evaluations should follow new-trial leadership for 7 years.

The same question arises for any fellowship that pays for a trial the fellow proposes. Because the WHO Trial Registration Data Set includes secondary identifying numbers and sources of monetary or material support among the items registries record [25], the split between proposed and new trials used here could be applied to fellowship schemes outside the United States wherever registrations list the award.

### Limitations

This study has limitations. First, trial plans come from funded abstracts, and cohorts of funded awardees cannot separate changes in behavior from changes in who was funded. Second, small validation cells constrain the correction, trial plans were treated as measured without error in the primary correction, and the raters were authors who saw award dates. Third, the pathway began on the same application date as NIH-wide changes to clinical trial applications, so no comparison separates them, and either change could have altered how often registrations cite the K grant, which classifies most K trials. Fourth, registry searches can miss unregistered trials, unnamed investigators, and name changes, and site roles fall outside the leadership definition. Fifth, award end dates were not retrieved, so new trials during and after the award could not be separated, and earlier support such as KL2 awards was not observed.

### Conclusions

In this cohort study, NIH’s dedicated K23 clinical trial pathway was associated with more awardees with trial plans leading their award-supported trial, although more complete registration or an earlier trend could not be ruled out. New-trial leadership within 5 years did not detectably increase, although gains of up to 11.5 percentage points remain possible under parallel trends. Whether the pathway produced more trial leaders is unclear. Evaluations of NIH’s 2027 KC1 award, which cannot fund trials, should track proposed and new trials separately.

## Author contributions

Conceptualization: Haining Wang, Hengyi Fu, Zuoyu Tian. Data curation: Haining Wang, Hengyi Fu, Zuoyu Tian. Formal analysis: Haining Wang, Zuoyu Tian. Investigation: Haining Wang. Methodology: Haining Wang, Zuoyu Tian. Software: Haining Wang. Supervision: Hengyi Fu. Validation: Haining Wang, Hengyi Fu, Zuoyu Tian. Visualization: Haining Wang. Writing – original draft: Haining Wang. Writing – review and editing: Haining Wang, Hengyi Fu, Zuoyu Tian.

## Supporting information

**S1 Appendix. Supplementary methods, tables, and figures.** Measurement instrument, human validation, and cohort flow; Tables A to L; Figs A and B.

**S1 Checklist. STROBE checklist.** Location of each STROBE item in the manuscript.

**S1 Appendix. Supplementary methods, tables, and figures**

Leadership of supported and new clinical trials after the NIH K23 clinical trial pathway

Terms used in this appendix: instrument = language model (uncorrected); regime = announcement type; stratum or period = early, legacy, or pathway awards; dossier = awardee record.

## Supplementary methods: measurement instrument, human validation, and cohort flow

**Candidate retrieval.** NIH RePORTER supplied the award cohorts, with trial outcomes retrieved from ClinicalTrials.gov by September 2026. Searches combined the awardee’s given name and surname, K grant identifiers, and surname with institution. Current registry records were supplemented by archived records from November 2022 and September 2024. The latter contributed responsible-party information only.

**Screening rules.** Trial start dates were assessed within the inclusive window from K start through five years later. Study type, K-grant linkage, and withdrawal with zero enrollment were determined by rule. Name matching required a surname with a compatible given name or initial within four tokens. Pairs failing the guard cannot qualify.

**Trial review.** Qwen/Qwen3.8-27B-FP8 was run locally on K abstracts and current and archived registry records to classify identity, leadership, intervention scope, and relationship to the K study. Protocol or published-protocol excerpts were also used in the contemporaneous analysis. All contemporaneous candidate pairs were reviewed. In the historical comparison, candidate pairs passing the name and timing screen were reviewed. The rest were classified as not qualifying by rule. The same registry sources and classification rules were used in both historical periods.

**Trial plans and validation.** Qwen/Qwen3.8-27B-FP8 classified K abstracts with funding-designation phrases masked as proposing an awardee-led interventional study, a mentor-led trial, other research, or unclear. The own-trial classification defined intent in the awarded, policy-responsive abstract. The leadership prompt was developed on held-out calibration pairs and frozen before the full run and human validation. Stored labels record prompt, schema and packet hashes. Weights and serving-container digests were not pinned, so new inference is a new measurement. Human validation design, agreement, instrument accuracy, and corrected estimates appear in Tables A, B, and D. Code, prompts and the rating codebook are available in the code repository.

**Table conventions**. Abbreviations are PI for principal investigator, DID for difference in differences, RD for risk difference, and CI for confidence interval. Unless otherwise stated, outcome windows extend from K start through five years later.

## Cohort-flow reconciliation

The contemporaneous and historical analyses use their own eligibility rules and the same follow-up requirements. The table distinguishes the separate baseline review used for the contemporaneous cohort from the common abstract-classifier exclusion used in both historical periods.

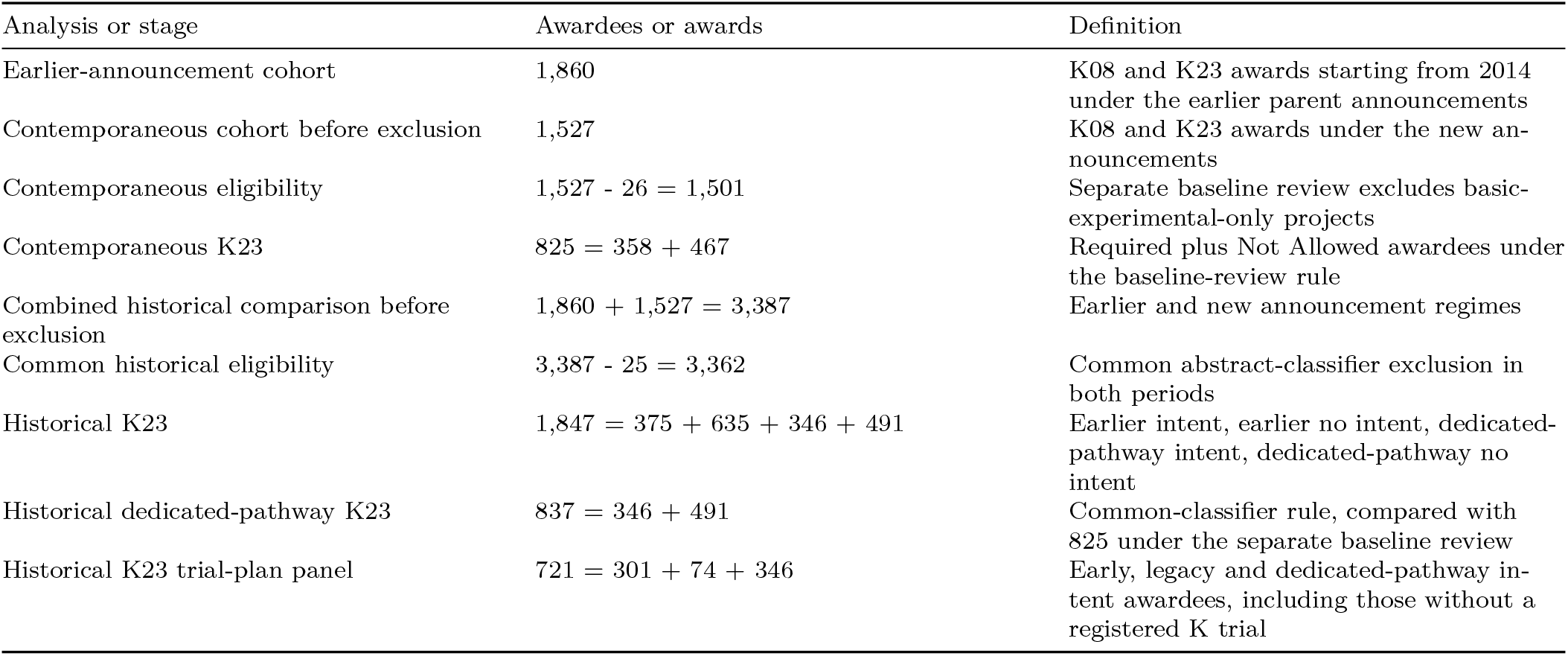

The 825 versus 837 counts reflect different exclusions, not different follow-up requirements. Legacy awards are identified by 2018–2019 award starts under PA-16 parent announcements. Their timing suggests likely dissemination-policy coverage, but application dates were not verified per award. Differences between displayed rates use unrounded estimates.

**Table A.**
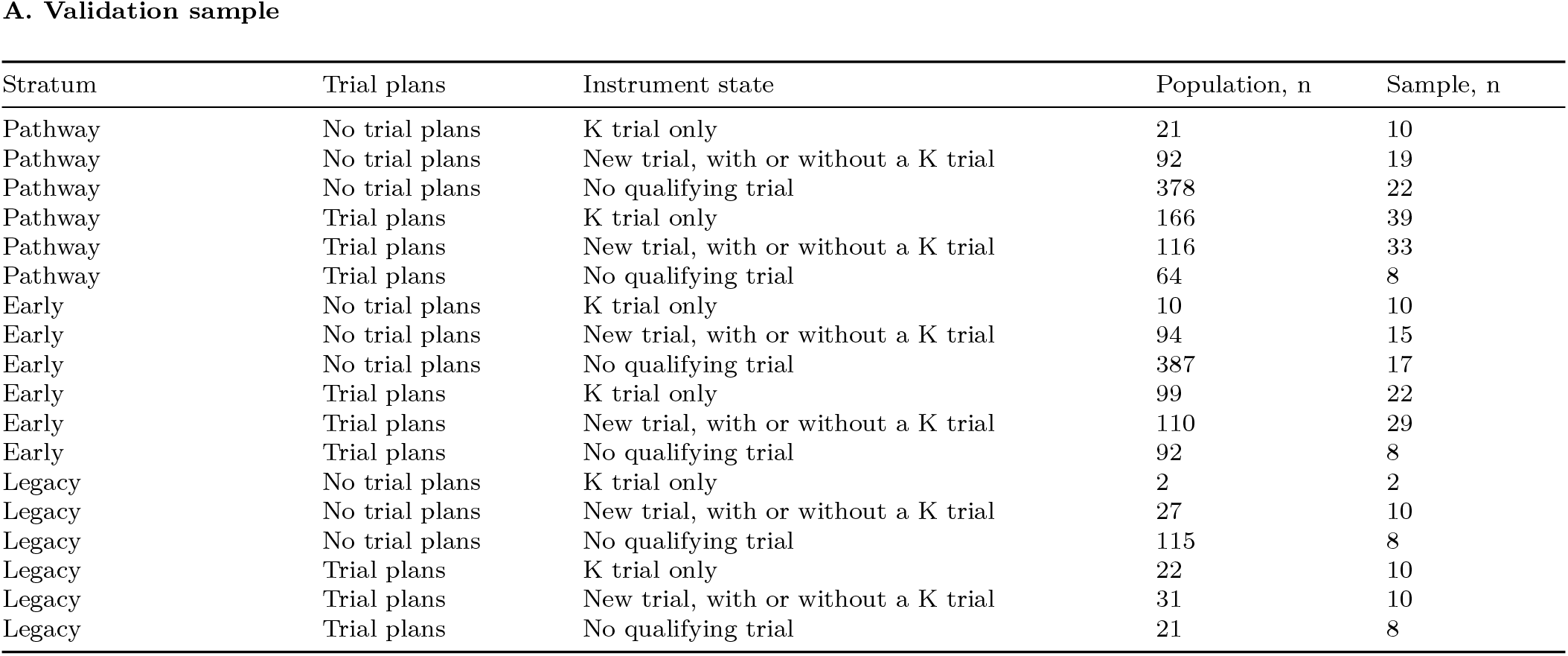
Human validation design. A stratified random sample of 280 historical K23 awardee dossiers represents 1,847 awardees across cells defined by stratum, baseline intent, and instrument state. Cell sizes targeted a 95% half-width of about 13 percentage points for the human-corrected K-trial DID. Two authors (HF and ZT) rated independently, blinded to announcement group, period labels, and model output, with ClinicalTrials.gov as the only trial-review source. Review included fresh name searches for missed trials, and funding-designation phrases were masked in award abstracts. Raters saw K start dates and project numbers needed for the five-year window and linkage, so timing and intent remained inferable. Of 110 double-rated dossiers, 70 used first-author resolution of disagreements and 40 used a reference rater randomly assigned before rating, while 170 were single-rated. A qualifying trial required an assigned intervention intended to prevent, diagnose, treat, or manage a condition or improve care, together with qualifying PI leadership. Measurement-only interventions did not meet this scope definition, even if the registry classified the study as interventional. K trials were grant-linked or judged to match the K plan, and new trials were unlinked and judged distinct, within five years of K start. The new-trial instrument state permits concurrent K-trial leadership. Six of 280 unclear awardee reference states were excluded from joint outcome correction cell counts, while the swapped-rater analysis excluded seven. Small cell sizes constrain precision, and human validation does not directly cover the additional seven-year observations. Intent is the instrument’s baseline own-trial classification. Cell populations and sample sizes sum to 1,847 and 280, respectively.

**Table B.**
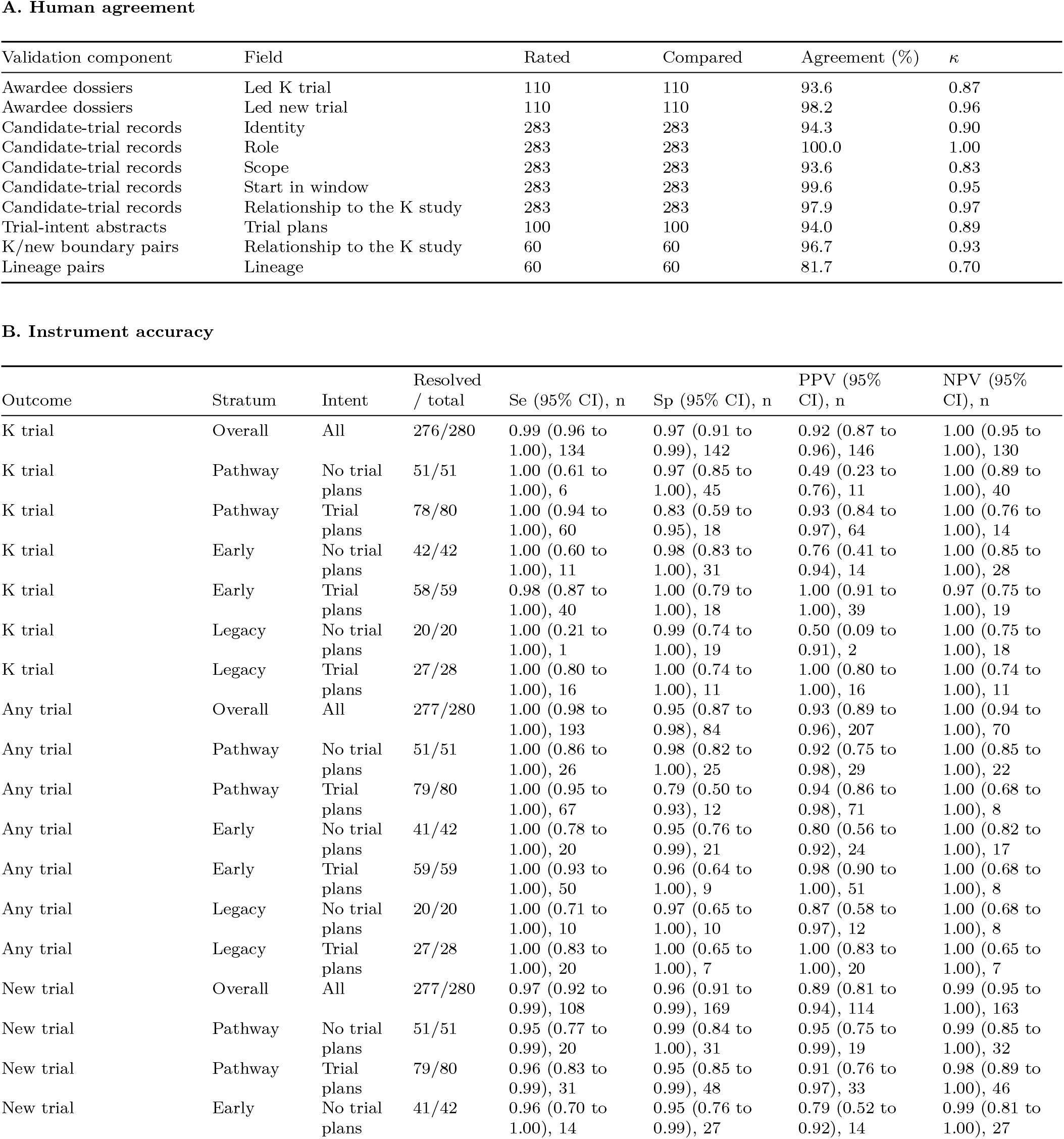
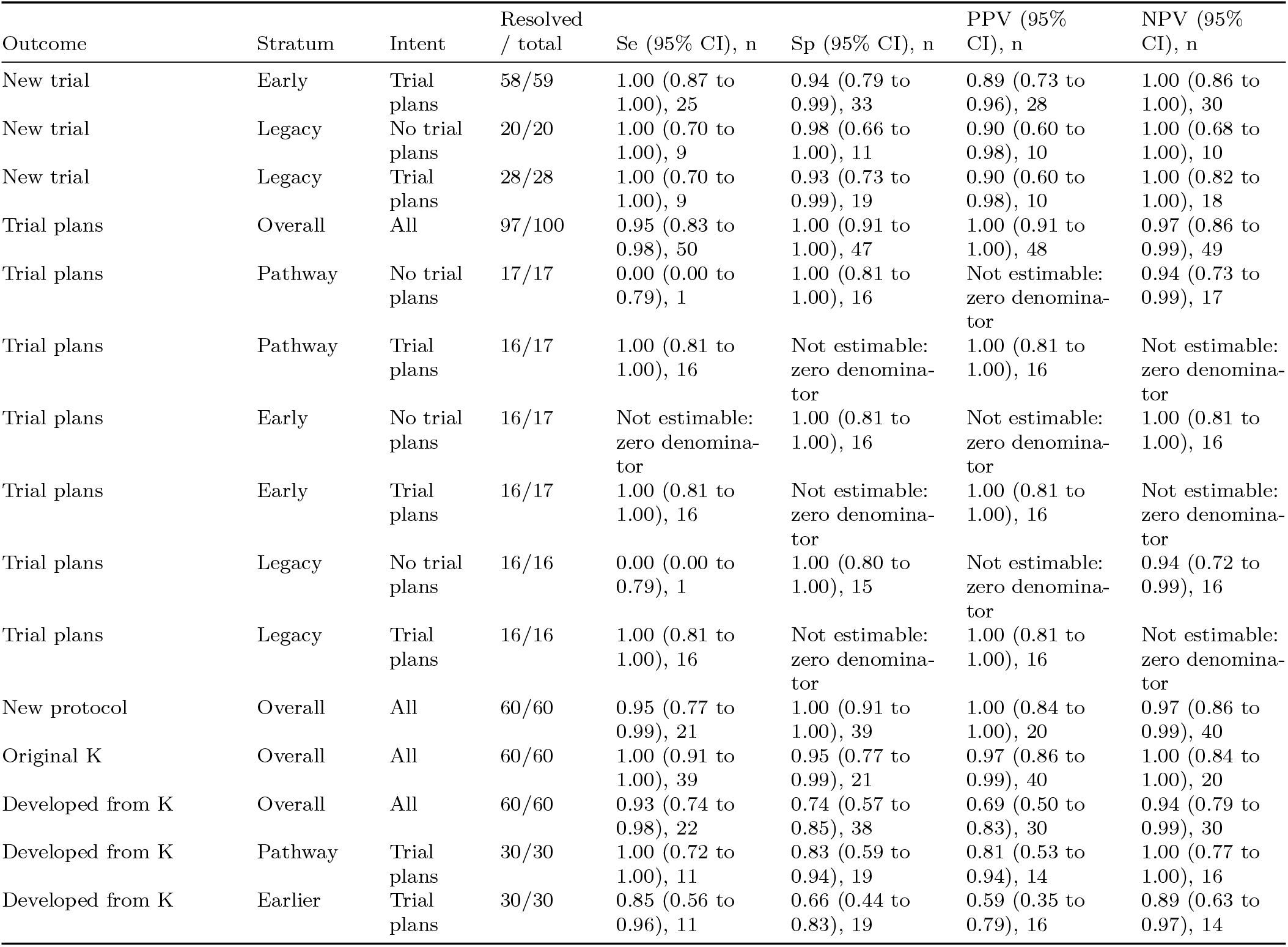
Human agreement and instrument accuracy. Asgreement covers all 110 double-rated dossiers and their 283 candidate-trial records, plus intent in 100 abstracts, K/new classification in 60 pairs, and lineage in 60 pairs. Agreement is between independent human raters, before reference-label resolution, and includes the full categorical fields. Accuracy compares instrument calls with the human reference, weighted by inverse sampling fractions, with Wilson intervals using Kish effective sample sizes. Outcome-specific unresolved references are excluded separately for accuracy, so its denominators differ from joint-state correction counts. Se is sensitivity, Sp specificity, PPV positive predictive value, and NPV negative predictive value, and n is the unweighted number of reviewed records in each denominator. Among awardees with trial plans, the model called a K trial for 4 of 18 reviewer-negative pathway records and 0 of 18 reviewer-negative early records. All leadership strata and intent groups are shown, together with stratum-specific intent, overall K/new accuracy and period-specific lineage accuracy. Reported measurement validity rests on this human sample. Qwen/Qwen3.8-27B-FP8 measured intent and person-trial status, and GPT-5.6 Sol rated lineage. Model lineage calls overstated continuity, especially in the earlier period. Agreement uses independent human ratings. Accuracy uses the corresponding human reference labels.

**Table C.**
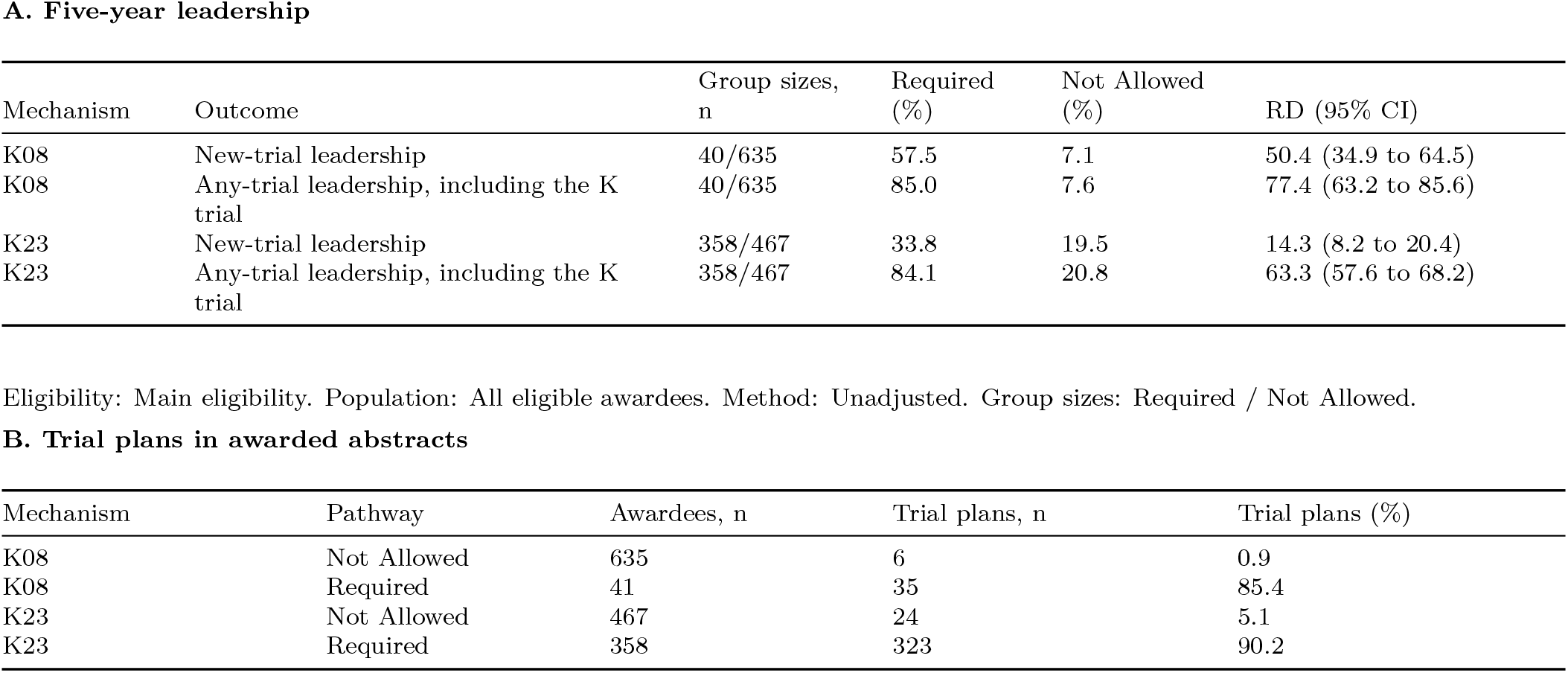
Contemporaneous Required versus Not Allowed comparison. Leadership within five years of K start among all eligible awardees. Risk differences and confidence limits are percentage points. The second panel gives the share of awarded abstracts classified as proposing the awardee’s own trial.

**Table D.**
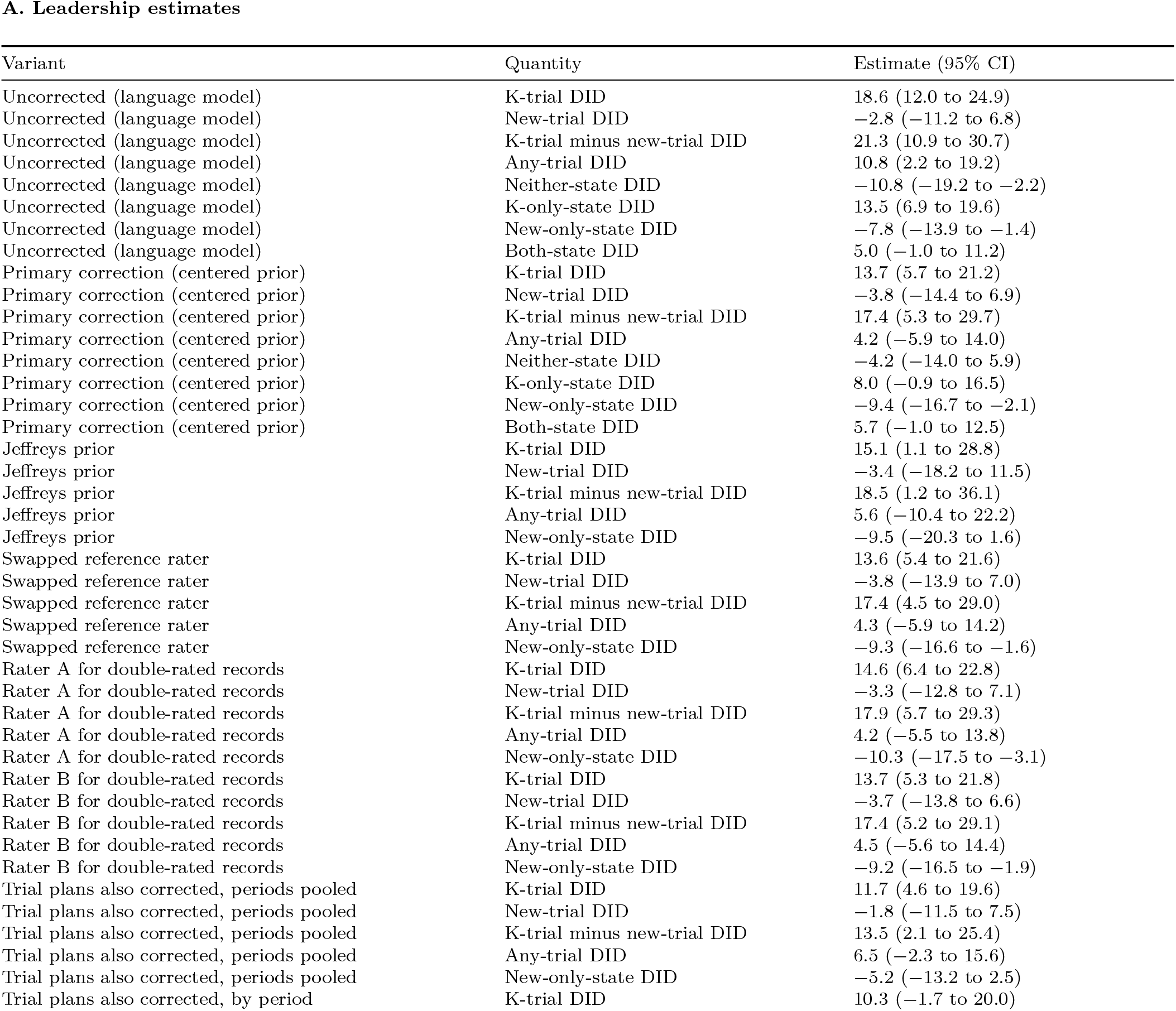
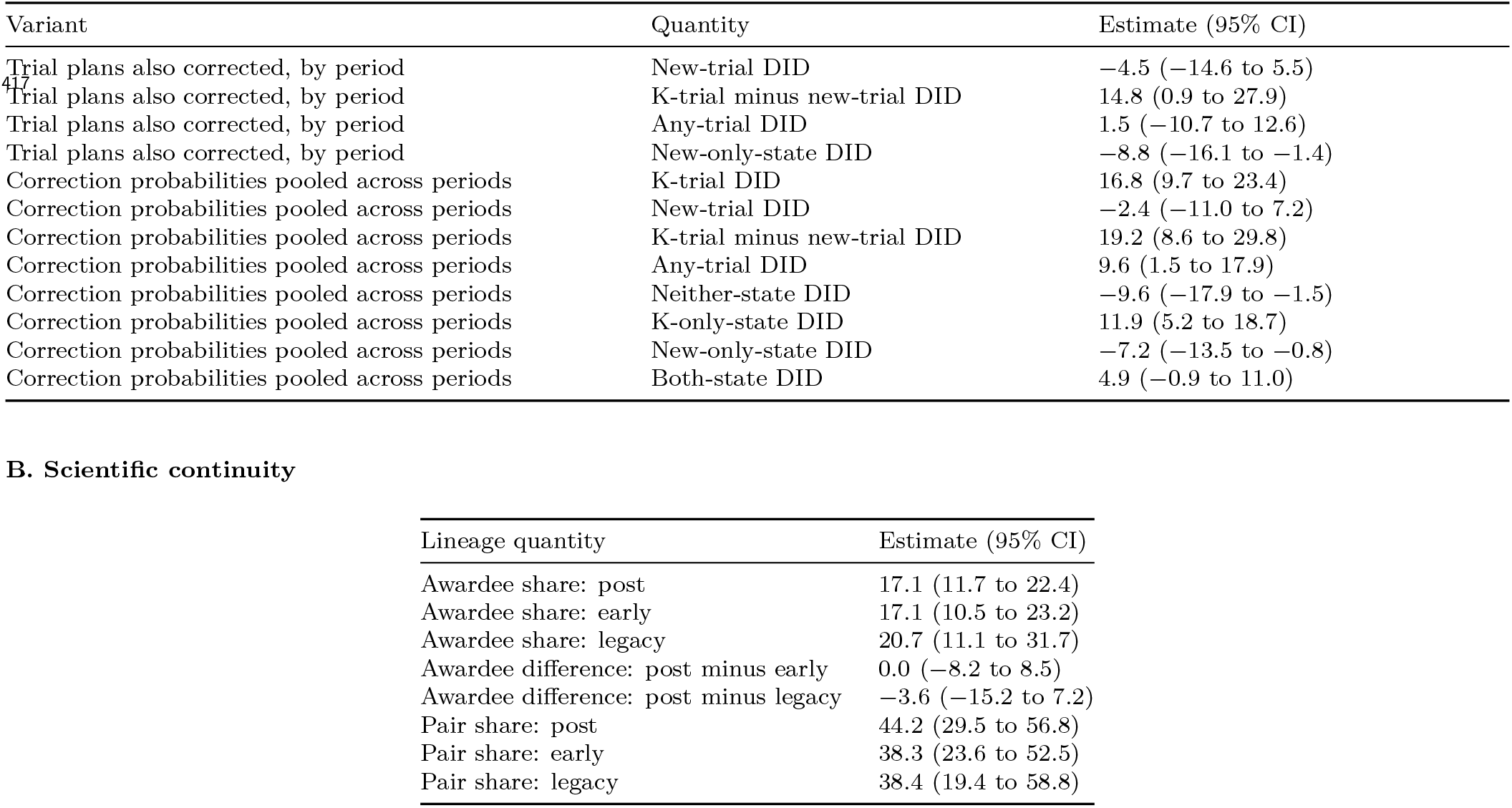
Instrument-measured and human-corrected estimates. DIDs use the same fiscal-year and institute-adjusted linear probability models in all 1,847 historical K23 awardees, with joint institution resampling for paired outcomes. Rated awardees take their resolved human joint state (neither, K only, new only, or both). Unrated or unresolved states are imputed within stratum x instrument intent x instrument joint-state cells. Empty refined cells borrow their three-state parent. The centered Dirichlet prior places 0.5 on the instrument state and 0.01 on each alternative. Jeffreys places 0.5 on each state. Swapping changes the assigned rater for the 40 overlap dossiers. Rater A and B variants additionally replace adjudication on the 70 initial double-rated dossiers. Single-rated dossiers keep their sole rating. Intent sensitivities use resolved human intent and impute other intents using either instrument call x instrument any-trial state (pooled across strata), or stratum x instrument call, with a centered Beta prior (0.5 on the call, 0.01 on the alternative). These are assumption-dependent reclassification sensitivities. For an unrated awardee in stratum s with instrument intent i and instrument joint state m, the human joint state is drawn from the Dirichlet posterior of P(human state | s, i, m). This assumes that rated and unrated awardees in a cell are exchangeable and that the distribution does not vary with fiscal year or institute within the cell. The pooled-period variant drops the stratum from the cells, so it assumes that P(human state | i, m) is the same in early, legacy, and dedicated-pathway awards. Reference counts remain fixed during institution resampling, so calibration uncertainty comes only from the Dirichlet posterior and does not reflect clustering of classification errors within institutions. All variants use 2,000 institution-bootstrap draws and percentile intervals. Corrected point estimates are draw means. Joint-state DIDs are shown for the instrument, the centered correction, and the pooled-period correction, and sum to zero before rounding. The new-only-state DID is shown for every variant. The two variants with the widest intervals reflect sparse cells (trial plans also corrected by period) and a flatter prior (Jeffreys), not estimates that point the other way. The instrument any-trial state here is the union of K and new trial leadership. Lineage correction draws binary developed-from-K statuses within period *×* K-trial availability *×* model-call cells, using a centered Beta prior inside 2,000 institution-bootstrap draws. The prior places 0.5 on the model call and 0.01 on the alternative. Awardee shares use all trial-plan awardees as denominators, while pair shares describe rated new-trial records. Units are percentage points for DIDs and differences and percentages for shares. Instrument estimates and human-corrected estimates use the outcome definitions and validation procedures described above.

**Table E.**
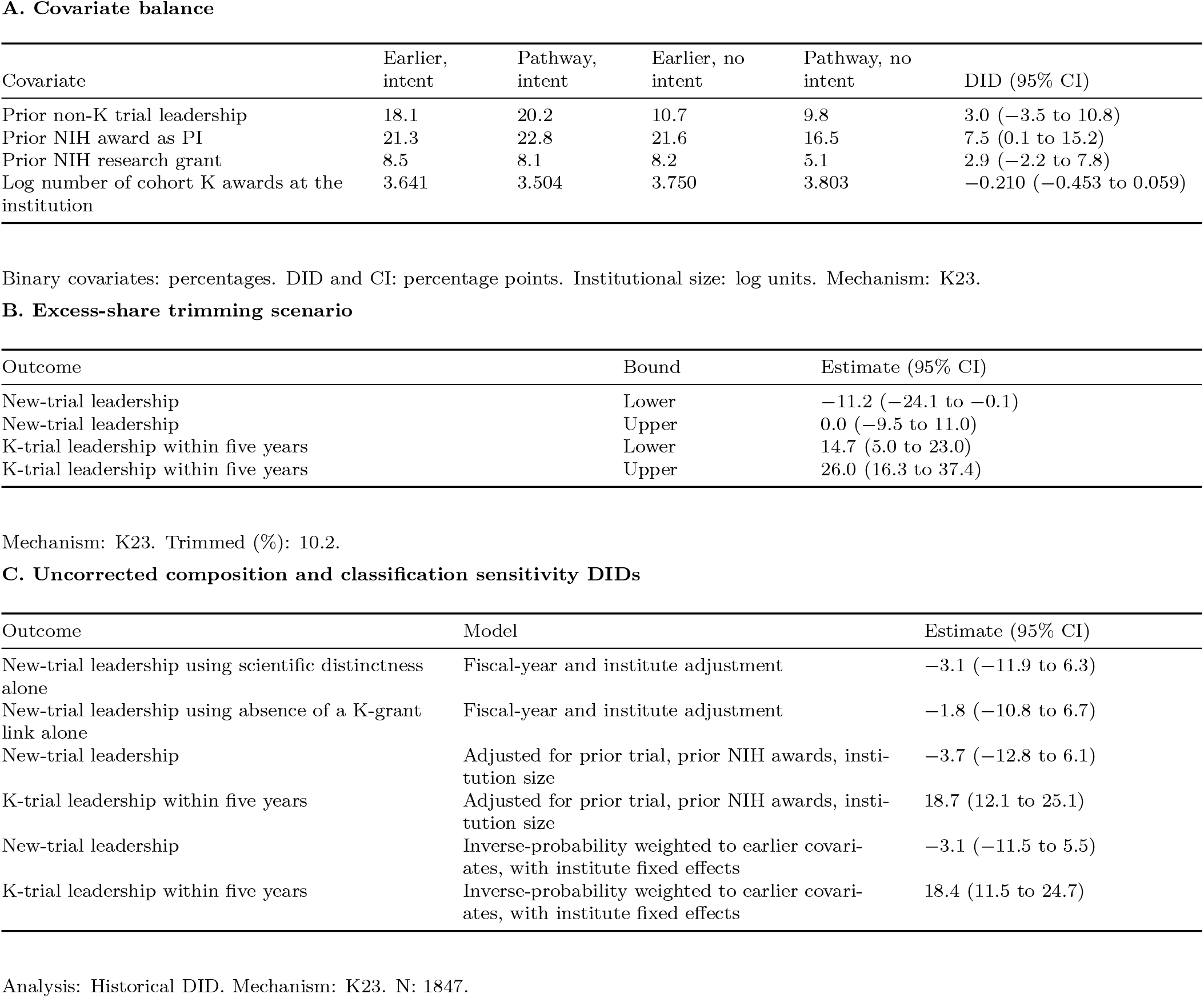
Composition checks. The first panel reports baseline covariates by trial plans and announcement regime, with the intent-by-period DID for each covariate. Binary covariate means are percentages, and covariate DIDs and confidence limits are percentage points. Institutional size is the log number of cohort K awards. The second panel removes the dedicated-pathway excess intent membership from the top or bottom of the outcome distribution. This trimming scenario assumes added membership without displacement and an unchanged comparator, so it does not bound arbitrary funding selection. Trimming bounds and their confidence limits are percentage points. The third panel uses uncorrected labels: covariate-adjusted and reweighted DIDs, and new-trial DIDs defined by scientific distinctness from the K plan alone (ignoring K-grant links) or by absence of a K-grant link alone.

**Table F.**
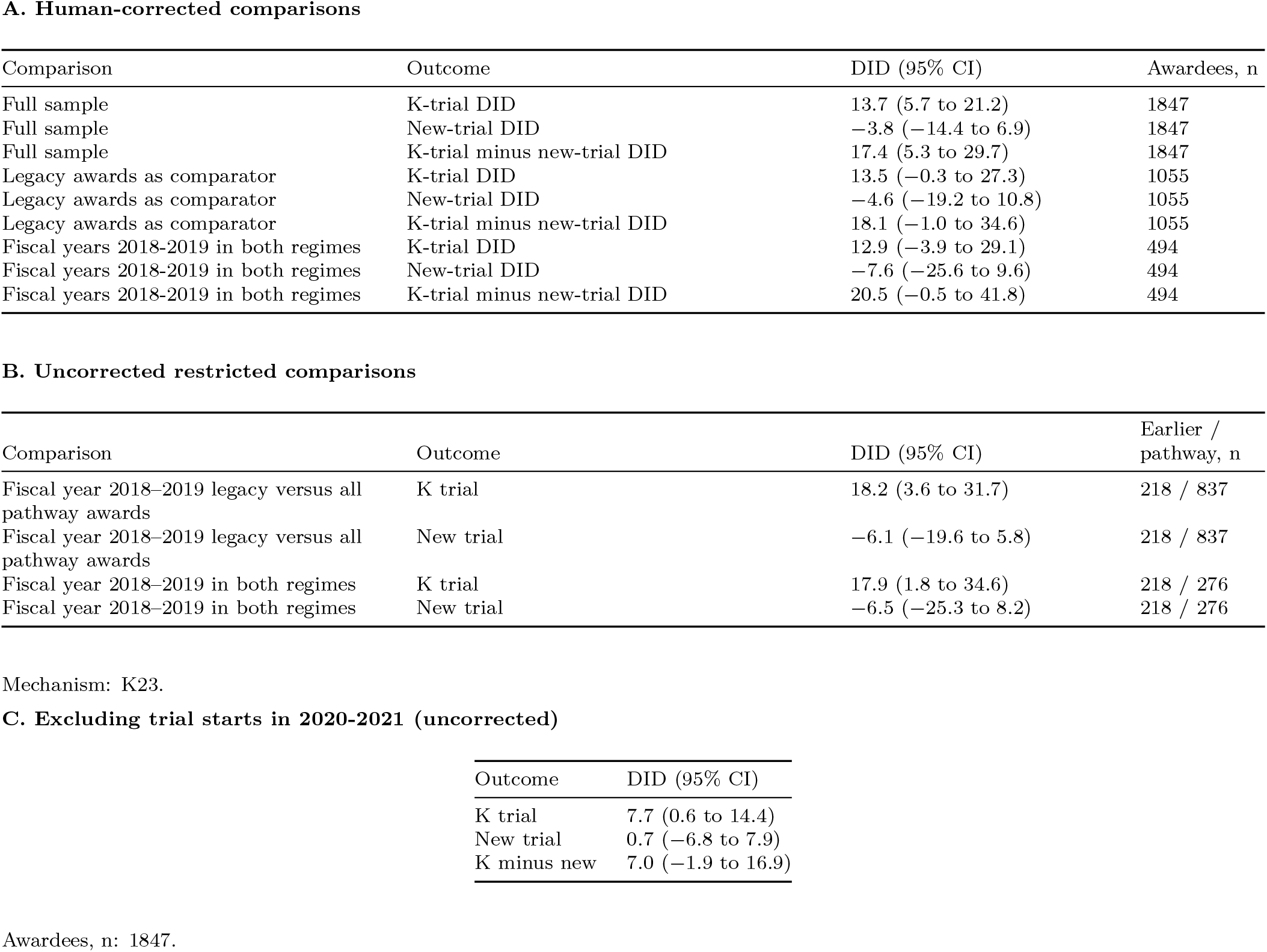
Comparison-group, calendar, and pandemic checks. All outcomes use five years from K start and fiscal-year and institute adjustment. The first panel is human corrected, computed in the same 2,000 institution-bootstrap draws as the primary estimates, so its full-sample rows equal the primary estimates. The comparison restricted to fiscal years 2018–2019 in both regimes aligns calendar time, including pandemic exposure, and consists mostly of fiscal year 2018 legacy and fiscal year 2019 dedicated-pathway awards. It does not separate policies that took effect for applications due from January 25, 2018, which applied to dedicated-pathway but not legacy applications, and it leaves selection into announcements unresolved. The second panel repeats the restricted comparisons with uncorrected (instrument) labels and 1,000 draws. The third panel, also uncorrected, drops trial starts in 2020–2021 while retaining all awardees. Because dedicated-pathway awardees started later, this exclusion removes more of their follow-up window than of earlier awardees’ windows and is biased toward smaller estimates.

**Table G.**
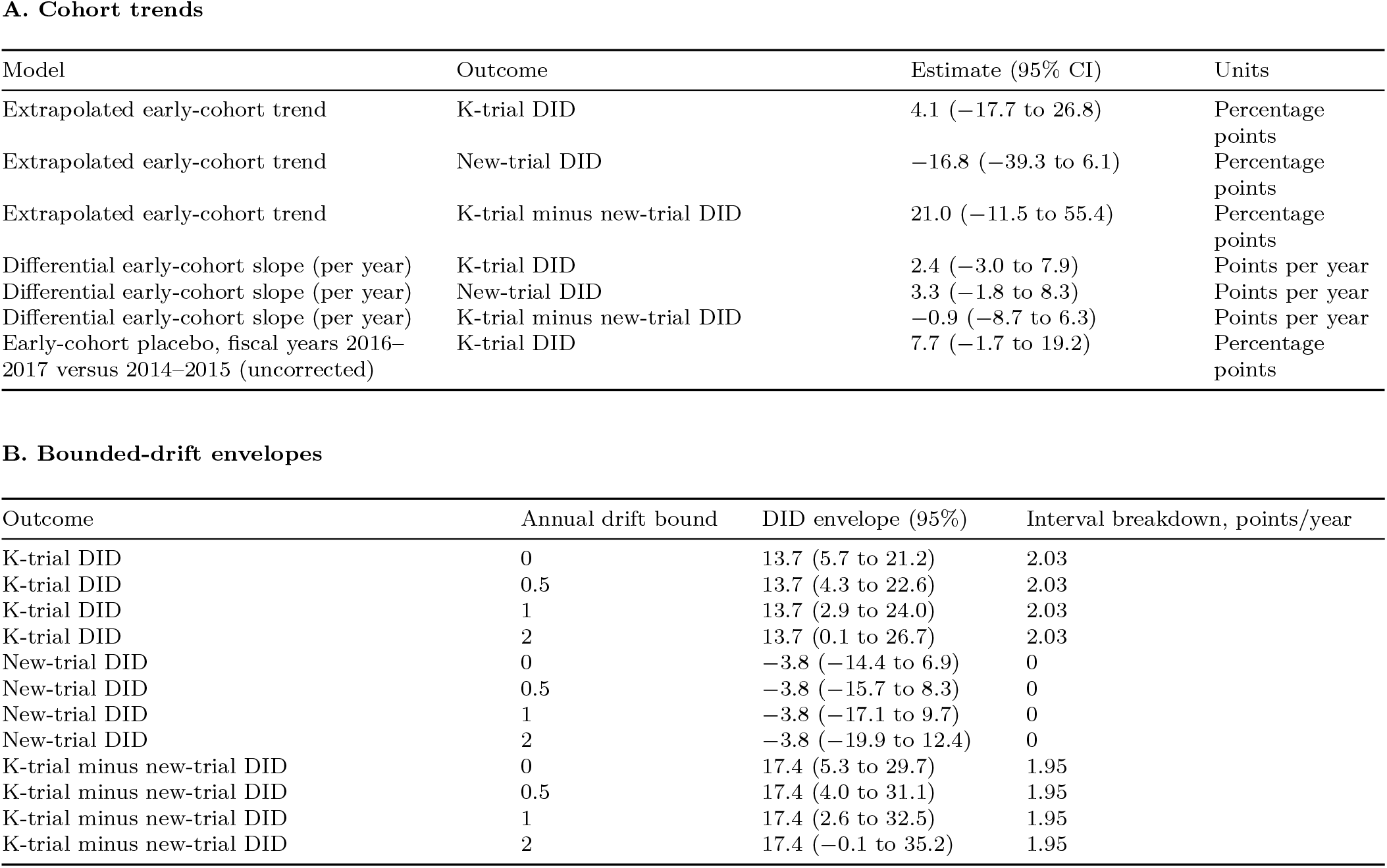
Cohort trends and bounded drift. Human corrected, in the primary 2,000 institution-bootstrap draws. Fiscal-year 2014–2017 intent-specific slopes adjust for year and institute. Subtracting this slope before refitting the DID gives the extrapolated estimate. The drift envelope permits additional intent-specific annual drift after fiscal year 2017, shifting each draw by its own drift exposure. Breakdown denotes the annual bound where the interval first includes zero. These additive-bias envelopes are not formal robust event-study confidence sets. Early-cohort follow-up extends into the policy era, precluding uncontaminated earlier outcome leads. The placebo compares the trial-plan gap in fiscal years 2016–2017 with 2014–2015 among early awards only. The mean drift exposure is 2.76 years, so the breakdown approximately equals the lower confidence limit divided by this exposure.

**Table H.**
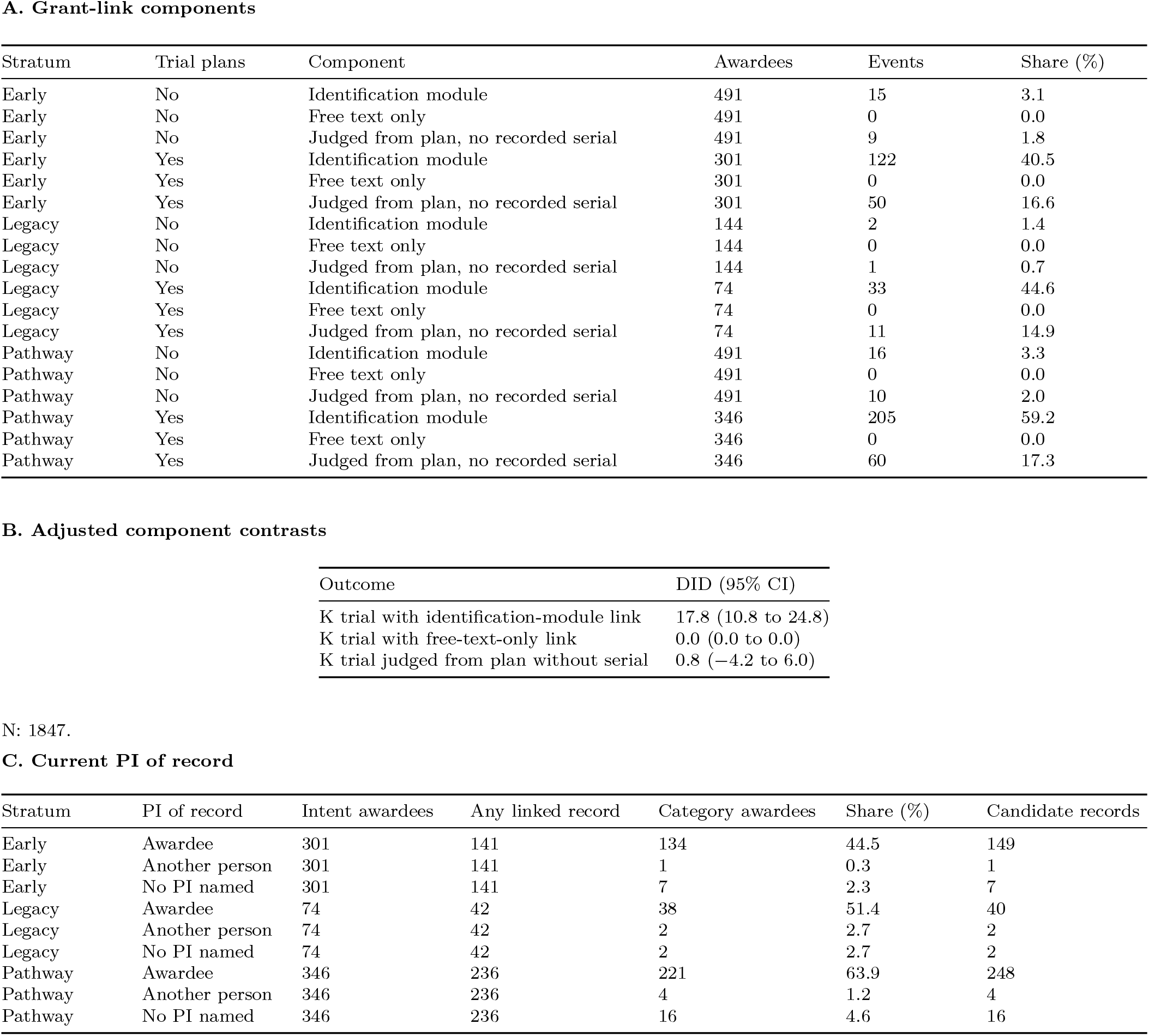
Grant-link decomposition and PI of record. Historical K-grant matching uses the identification module, with no free-text-only qualification. No recorded serial means no identification-module link, rather than a search proving its absence throughout free text. Component awardee shares can overlap across trials, so component DIDs need not sum to the total. PI diagnostics use current fields on all linked candidate records, including observational records and records outside the outcome window. Awardee PI takes precedence when co-listed with another person. Record categories are exclusive, while awardee categories may overlap. Shares use all intent awardees as denominators and do not establish initial PI designation or actual responsibility. Component contrasts use fiscal-year and institute adjustment with 1,000 institution draws.

**Table I.**
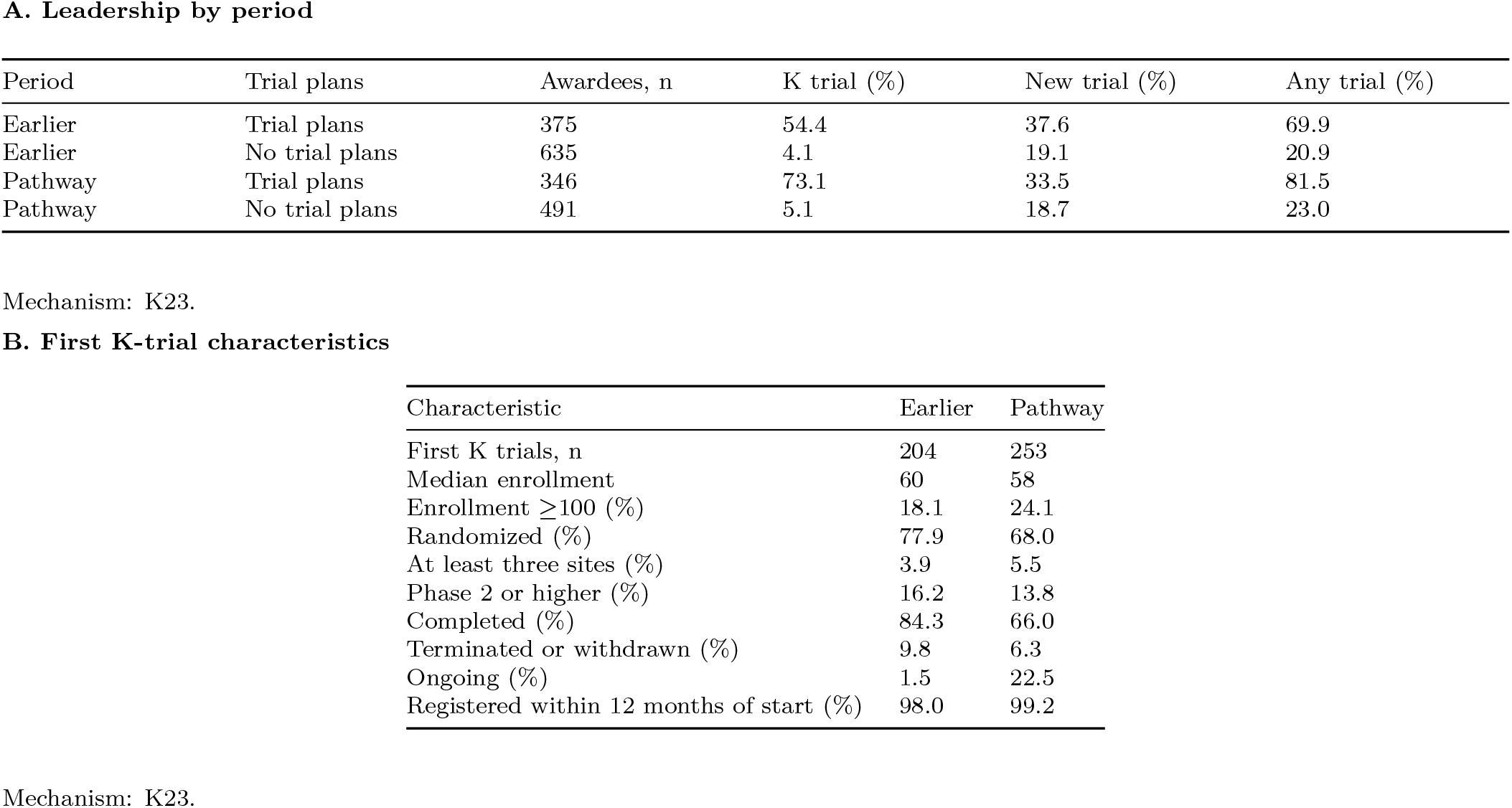
Leadership by period and trial plans. Instrument-measured leadership within five years of K start. Percentages count awardees. K trials either cite the K grant or are judged to match the K plan. The second panel describes the first K trial starting within five years of K start for each trial-plan awardee with such a trial, by period. Results reporting is in Table J.

**Table J.**
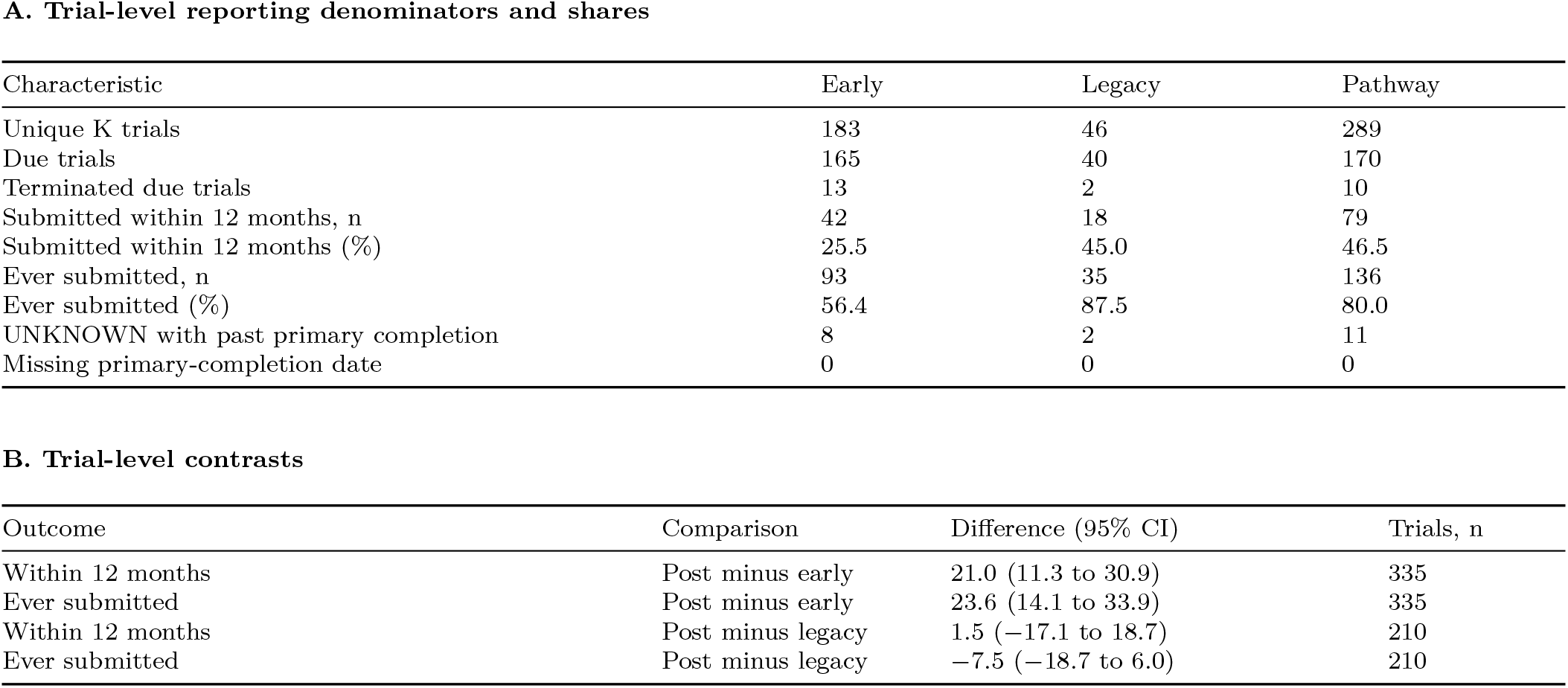
Trial-level reporting. The reported outcome is trial-level first results submission within twelve months of primary completion. Due trials were COMPLETED or TERMINATED with positive ACTUAL enrollment, with primary completion at least twelve calendar months before September 27, 2026. Month-only dates use the first day. Primary-completion ACTUAL/ESTIMATED type was not retrieved and was not imposed. UNKNOWN trials with past completion dates are reported separately and excluded from due denominators. Each eligible trial appears once, with no duplicates across awardees or strata. Both trial metrics use first results submission, not public posting after quality control. Ever submission includes reports after twelve months and remains subject to unequal follow-up. All intervals use institution resampling.

**Table K.**
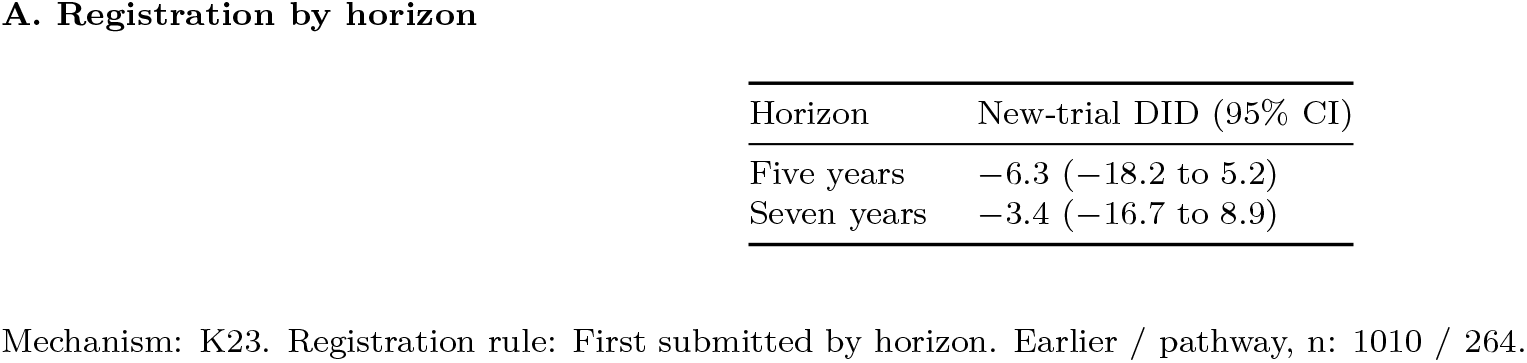
Registration by the follow-up horizon. Both horizons use the same complete-seven-year K23 sample. Symmetric registration requires first submission no later than K start plus the respective horizon. Five-year outcomes are nested within seven-year outcomes, with paired institution draws and fiscal-year and institute adjustment. The rule excludes retrospective registrations after each horizon but cannot recover never-registered or unretrieved trials. Human leadership validation covers five years only.

**Table L.**
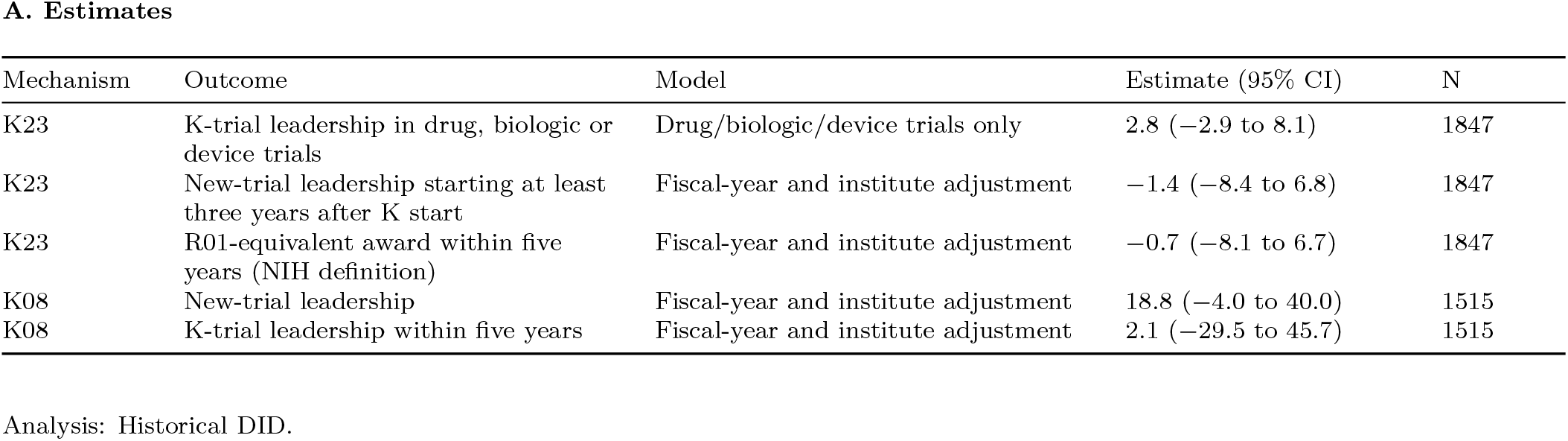
Further historical estimates. Historical DIDs adjust for fiscal year and institute. Estimates and confidence limits are percentage points. The drug, biologic or device restriction counts only K trials testing those interventions. Late-start new trials begin at least three years after K start. R01-equivalent awards within five years follow the NIH Data Book definition: DP1, DP2, DP5, R01, R37, R56, RF1, RL1 and U01, plus R35 from NIGMS or NHGRI only. K08 estimates rest on 14 earlier trial-plan awardees.

**Fig A.**
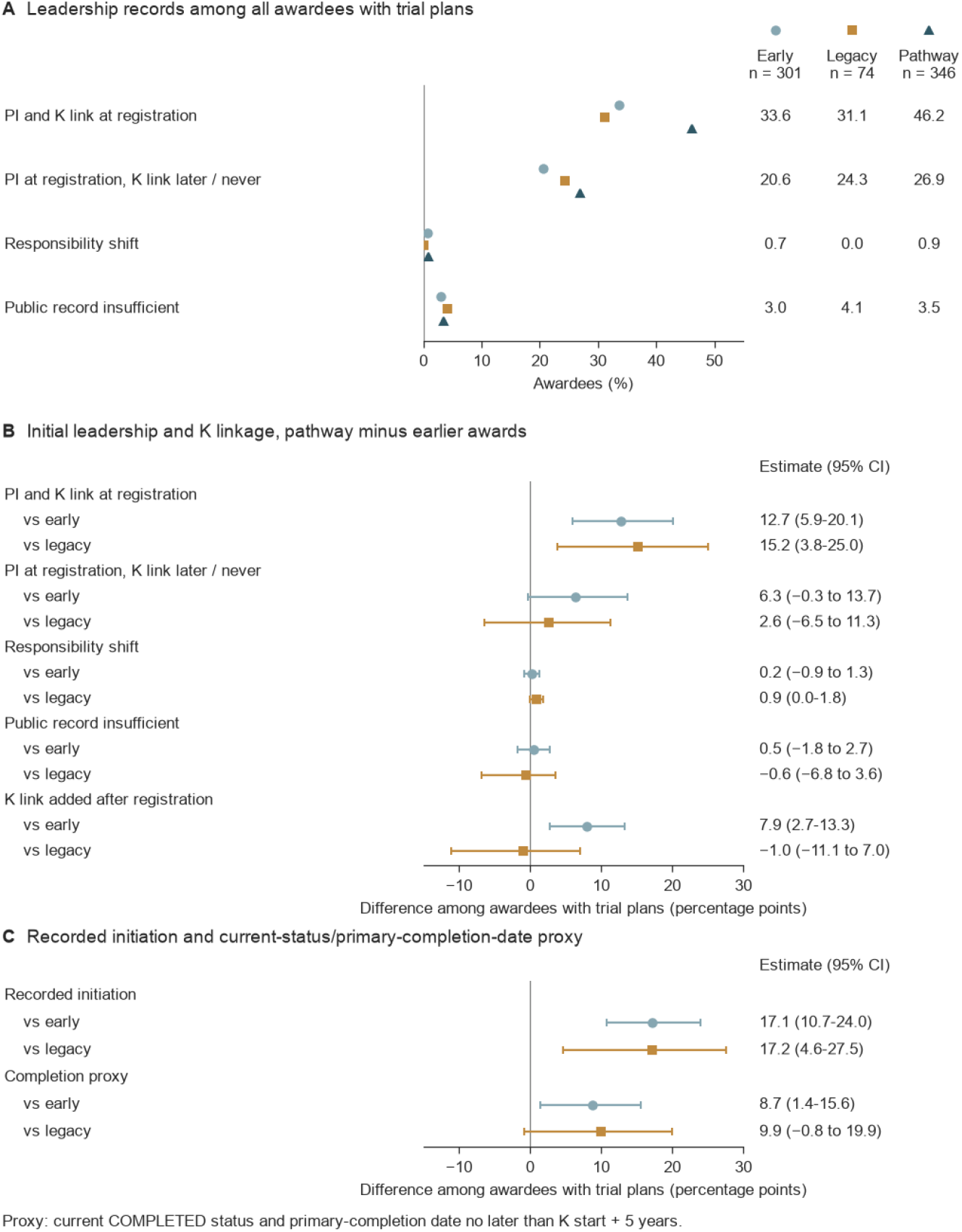
Registration categories, initial leadership, and recorded initiation among K23 awardees with trial plans. A, Registration categories are exclusive for individual trials, although awardees can contribute trials to several categories. B, Initial leadership and K linkage differences between dedicated-pathway awards and early or legacy awards. C, Recorded initiation and a current-status/primary-completion-date proxy, with all trial-plan awardees as denominators. Recorded initiation means that a K trial was recruiting, enrolling by invitation, active, or completed in any registry version, or reported actual enrollment above zero. It was documented for 52.8%, 52.7%, and 69.9% of early, legacy, and pathway awardees with trial plans, and the completion proxy for 30.9%, 29.7%, and 39.6%. Completion requires current COMPLETED status and primary completion within 5 years of K start, without verifying overall completion. Whiskers show 95% CIs from bootstrap draws resampling institutions. Shares and differences are uncorrected and compare awardees with trial plans across strata. They are not difference-in-differences estimates.

**Fig B.**
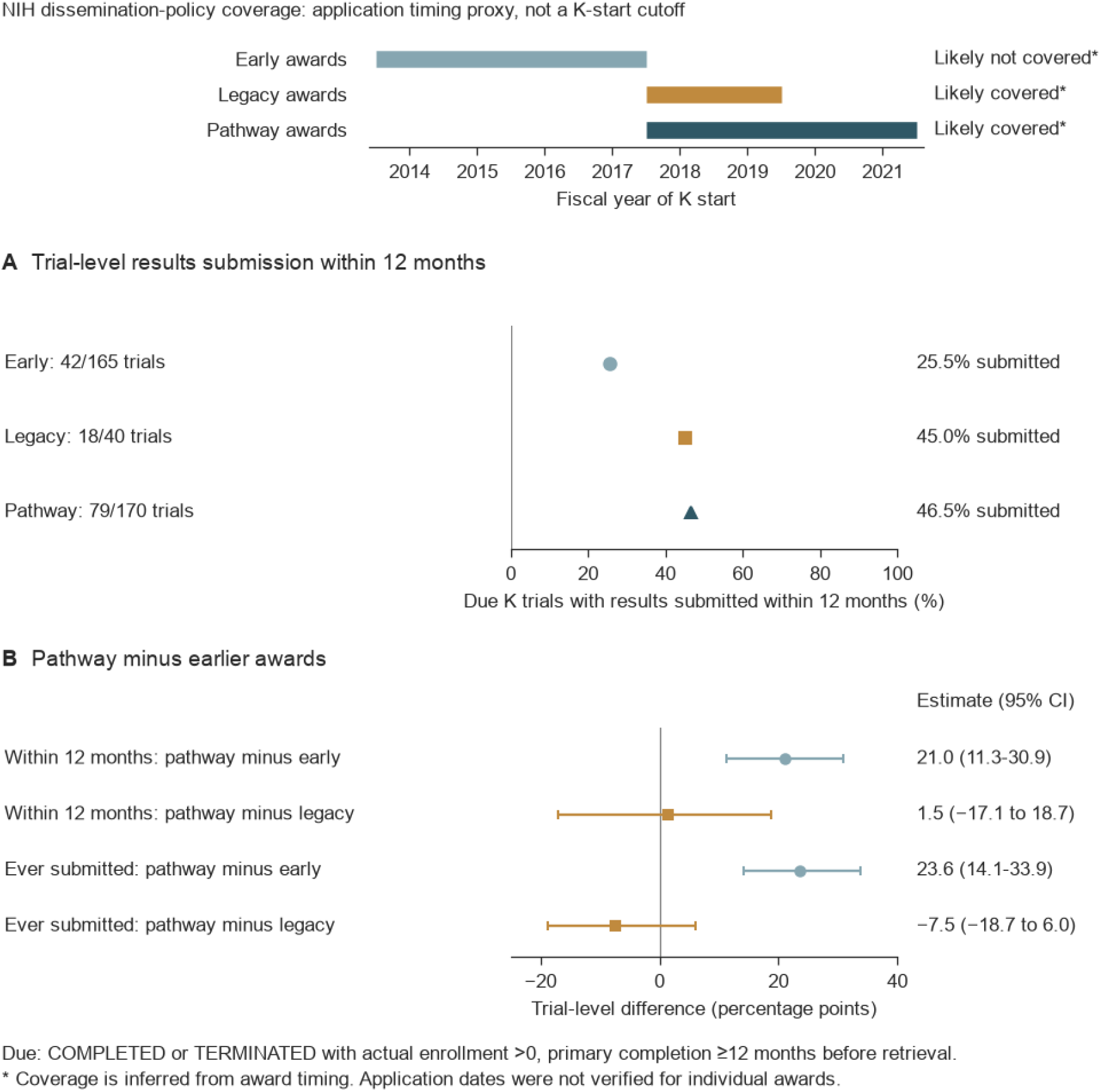
Trial-level results reporting by award stratum. The timeline shows fiscal years of K start. Coverage labels summarize a timing proxy for the NIH dissemination policy, and individual application dates were not verified. A, Results submission within 12 months of primary completion among due trials, which were COMPLETED or TERMINATED with positive actual enrollment and primary completion at least 12 months before retrieval. B, Differences between dedicated-pathway and earlier strata in within-12-month and ever-submitted results. Whiskers show 95% CIs from bootstrap draws resampling institutions.

## References

[1] Garrison HH, Deschamps AM. NIH research funding and early career physician scientists: continuing challenges in the 21st century. FASEB J. 2014;28(3):1049–1058. doi:10.1096/fj.13-241687.

[2] National Institutes of Health. NOT-OD-26-098: Notice of NIH updates to career development (K) award programs; 2026. NIH Guide for Grants and Contracts; released August 12, 2026 [cited 2026 Sep 28]. Available from: https://grants.nih.gov/grants/guide/notice-files/NOT-OD-26-098.html.

[3] Brandt PD, Sturzenegger Varvayanis S, Baas T, Bolgioni AF, Alder J, Petrie KA, et al. A cross-institutional analysis of the effects of broadening trainee professional development on research productivity. PLoS Biol. 2021;19(7):e3000956. doi:10.1371/journal.pbio.3000956.

[4] Organisation for Economic Co-operation and Development. Government R&D funding and company behaviour: measuring behavioural additionality. Paris: OECD Publishing; 2006. Available from: 10.1787/9789264025851-en.

[5] Jacob BA, Lefgren L. The impact of research grant funding on scientific productivity. J Public Econ. 2011;95(9-10):1168–1177. doi:10.1016/j.jpubeco.2011.05.005.

[6] Nikaj S, Lund PK. The impact of individual mentored career development (K) awards on the research trajectories of early-career scientists. Acad Med. 2019;94(5):708–714. doi:10.1097/ACM.0000000000002543.

[7] Li D, Agha L. Big names or big ideas: do peer-review panels select the best science proposals? Science. 2015;348(6233):434–438. doi:10.1126/science.aaa0185.

[8] National Institutes of Health. NOT-OD-17-043: Update on clinical trial funding opportunity announcement policy; 2017. NIH Guide for Grants and Contracts; released February 17, 2017 [cited 2026 Oct 4]. Available from: https://grants.nih.gov/grants/guide/notice-files/NOT-OD-17-043.html.

[9] National Institutes of Health. PA-16-198: Mentored Patient-Oriented Research Career Development Award (Parent K23); 2016. NIH Guide for Grants and Contracts; posted April 15, 2016 [cited 2026 Sep 28]. Available from: https://grants.nih.gov/grants/guide/pa-files/PA-16-198.html.

[10] National Institutes of Health. PA-16-191: Mentored Clinical Scientist Research Career Development Award (Parent K08); 2016. NIH Guide for Grants and Contracts; posted April 15, 2016 [cited 2026 Sep 28]. Available from: https://grants.nih.gov/grants/guide/pa-files/PA-16-191.html.

[11] National Institutes of Health. NOT-OD-18-001: Clinical trials: special considerations for career development, fellowship, training, and research education programs; 2017. NIH Guide for Grants and Contracts; released November 6, 2017 [cited 2026 Sep 28]. Available from: https://grants.nih.gov/grants/guide/notice-files/NOT-OD-18-001.html.

[12] National Institutes of Health. Table of IC-specific information, requirements and staff contacts: parent K23, independent clinical trial required; 2026. NIH Grants and Funding [cited 2026 Oct 16]. Available from: https://grants.nih.gov/grants/guide/contacts/parent-K23-CT-required.html.

[13] Baldwin TT, Ford JK. Transfer of training: a review and directions for future research. Pers Psychol. 1988;41(1):63–105. doi:10.1111/j.1744-6570.1988.tb00632.x.

[14] Axfors C, Nilsonne G. Nordic trial reporting project: raw data from EU Clinical Trials Registry (EUCTR) and ClinicalTrials.gov [data set, including the AACT database of November 9, 2022]; 2023. Zenodo; published November 9, 2023. Available from: 10.5281/zenodo.10091147.

[15] Clinical Trials Transformation Initiative. Clinical Trials Transformation Initiative’s (CTTI) Trial Information through September 2024 [data set]; 2024. Zenodo; published October 23, 2024. Available from: 10.5281/zenodo.13984069.

[16] National Institutes of Health. NIH Data Book: R01-equivalent grants; 2026. NIH Re-PORT; updated March 2026 [cited 2026 Sep 29]. Available from: https://report.nih.gov/nihdatabook/category/3.

[17] Cole SR, Chu H, Greenland S. Multiple-imputation for measurement-error correction. Int J Epidemiol. 2006;35(4):1074–1081. doi:10.1093/ije/dyl097.

[18] Derington CG, Cervantes L, Allen LA, Trinkley KE, Huebschmann AG, Ho PM. Clinical trial components of NIH career development awards: supporting the nation’s future trialists. medRxiv [preprint]. 2026;doi:10.64898/2026.09.18.26363419.

[19] Swartz HA. Career development awards for clinical trialists: a training pathway under threat. Am J Psychother. 2026;79(3):105–106. doi:10.1176/appi.psychotherapy.26079001.

[20] Boulanger-Espeut C, Demblowski L, Schmidt M, Torres S, Bogler O, Zahir N. Outcome evaluation of NCI Clinical Scientist Research Career Development Award (K08) program. J Cancer Educ. 2026;doi:10.1007/s13187-026-02896-z.

[21] National Institutes of Health. NOT-OD-15-015: Notice of revised NIH definition of clinical trial; 2014. NIH Guide for Grants and Contracts; released October 23, 2014 [cited 2026 Oct 4]. Available from: https://grants.nih.gov/grants/guide/notice-files/NOT-OD-15-015.html.

[22] National Institutes of Health. NOT-OD-17-062: New “FORMS-E” grant application forms and instructions coming for due dates on or after January 25, 2018; 2017. NIH Guide for Grants and Contracts; released April 27, 2017 [cited 2026 Oct 4]. Available from: https://grants.nih.gov/grants/guide/notice-files/NOT-OD-17-062.html.

[23] US Department of Health and Human Services. Clinical trials registration and results information submission: final rule (42 CFR Part 11). Fed Regist. 2016;81(183):64981–65157.

[24] National Institutes of Health. NOT-OD-16-149: NIH policy on the dissemination of NIH-funded clinical trial information; 2016. NIH Guide for Grants and Contracts; released September 16, 2016; effective January 18, 2017. Available from: https://grants.nih.gov/grants/guide/notice-files/NOT-OD-16-149.html.

[25] World Health Organization. WHO Trial Registration Data Set (version 1.3.1); 2026. International Clinical Trials Registry Platform [cited 2026 Oct 16]. Available from: https://www.who.int/tools/clinical-trials-registry-platform/network/who-data-set.

