## Supplemental Materials for "Leadership of supported and new clinical trials after the NIH K23 clinical trial pathway"

| Analysis or stage | Awardees or awards | Definition |
|---|---|---|
| Earlier-announcement cohort | 1,860 | K08 and K23 awards starting from 2014 under the earlier parent announcements |
| Contemporaneous cohort before exclusion | 1,527 | K08 and K23 awards under the new announcements |
| Contemporaneous eligibility | 1,527 - 26 = 1,501 | Separate baseline review excludes basic-experimental-only projects |
| Contemporaneous K23 | 825 = 358 + 467 | Required plus Not Allowed awardees under the baseline-review rule |
| Combined historical comparison before exclusion | 1,860 + 1,527 = 3,387 | Earlier and new announcement regimes |
| Common historical eligibility | 3,387 - 25 = 3,362 | Common abstract-classifier exclusion in both periods |
| Historical K23 | 1,847 = 375 + 635 + 346 + 491 | Earlier intent, earlier no intent, dedicated-pathway intent, dedicated-pathway no intent |
| Historical dedicated-pathway K23 | 837 = 346 + 491 | Common-classifier rule, compared with 825 under the separate baseline review |
| Historical K23 trial-plan panel | 721 = 301 + 74 + 346 | Early, legacy and dedicated-pathway intent awardees, including those without a registered K trial |

Intent is the instrument's baseline own-trial classification. Cell populations and sample sizes sum to 1,847 and 280, respectively.

**A. Validation sample**

| Stratum | Trial plans | Instrument state | Population, n | Sample, n |
|---|---|---|---|---|
| Pathway | No trial plans | K trial only | 21 | 10 |
| Pathway | No trial plans | New trial, with or without a K trial | 92 | 19 |
| Pathway | No trial plans | No qualifying trial | 378 | 22 |
| Pathway | Trial plans | K trial only | 166 | 39 |
| Pathway | Trial plans | New trial, with or without a K trial | 116 | 33 |
| Pathway | Trial plans | No qualifying trial | 64 | 8 |
| Early | No trial plans | K trial only | 10 | 10 |
| Early | No trial plans | New trial, with or without a K trial | 94 | 15 |
| Early | No trial plans | No qualifying trial | 387 | 17 |
| Early | Trial plans | K trial only | 99 | 22 |
| Early | Trial plans | New trial, with or without a K trial | 110 | 29 |
| Early | Trial plans | No qualifying trial | 92 | 8 |
| Legacy | No trial plans | K trial only | 2 | 2 |
| Legacy | No trial plans | New trial, with or without a K trial | 27 | 10 |
| Legacy | No trial plans | No qualifying trial | 115 | 8 |
| Legacy | Trial plans | K trial only | 22 | 10 |
| Legacy | Trial plans | New trial, with or without a K trial | 31 | 10 |
| Legacy | Trial plans | No qualifying trial | 21 | 8 |

### Table B. Human agreement and instrument accuracy

Agreement covers all 110 double-rated dossiers and their 283 candidate-trial records, plus intent in 100 abstracts, K/new classification in 60 pairs, and lineage in 60 pairs. Agreement is between independent human raters, before reference-label resolution, and includes the full categorical fields. Accuracy compares instrument calls with the human reference, weighted by inverse sampling fractions, with Wilson intervals using Kish effective sample sizes. Outcome-specific unresolved references are excluded separately for accuracy, so its denominators differ from joint-state correction counts. Se is sensitivity, Sp specificity, PPV positive predictive value, and NPV negative predictive value, and n is the unweighted number of reviewed records in each denominator. Among awardees with trial plans, the model called a K trial for 4 of 18 reviewer-negative pathway records and 0 of 18 reviewer-negative early records. All leadership strata and intent groups are shown, together with stratum-specific intent, overall K/new accuracy and period-specific lineage accuracy.

Reported measurement validity rests on this human sample. Qwen/Qwen3.8-27B-FP8 measured intent and person-trial status, and GPT-5.6 Sol rated lineage. Model lineage calls overstated continuity, especially in the earlier period.

Agreement uses independent human ratings. Accuracy uses the corresponding human reference labels.

**A. Human agreement**

| Validation component | Field | Rated | Compared | Agreement (%) | $\kappa$ |
|---|---|---|---|---|---|
| Awardee dossiers | Led K trial | 110 | 110 | 93.6 | 0.87 |
| Awardee dossiers | Led new trial | 110 | 110 | 98.2 | 0.96 |
| Candidate-trial records | Identity | 283 | 283 | 94.3 | 0.90 |
| Candidate-trial records | Role | 283 | 283 | 100.0 | 1.00 |
| Candidate-trial records | Scope | 283 | 283 | 93.6 | 0.83 |
| Candidate-trial records | Start in window | 283 | 283 | 99.6 | 0.95 |
| Candidate-trial records | Relationship to the K study | 283 | 283 | 97.9 | 0.97 |
| Trial-intent abstracts | Trial plans | 100 | 100 | 94.0 | 0.89 |
| K/new boundary pairs | Relationship to the K study | 60 | 60 | 96.7 | 0.93 |
| Lineage pairs | Lineage | 60 | 60 | 81.7 | 0.70 |

**B. Instrument accuracy**

| Outcome | Stratum | Intent | Resolved / total | Se (95% CI), n | Sp (95% CI), n | PPV (95% CI), n | NPV (95% CI), n |
|---|---|---|---|---|---|---|---|
| K trial | Overall | All | 276/280 | 0.99 (0.96 to 1.00), 134 | 0.97 (0.91 to 0.99), 142 | 0.92 (0.87 to 0.96), 146 | 1.00 (0.95 to 1.00), 130 |
| K trial | Pathway | No trial plans | 51/51 | 1.00 (0.61 to 1.00), 6 | 0.97 (0.85 to 1.00), 45 | 0.49 (0.23 to 0.76), 11 | 1.00 (0.89 to 1.00), 40 |
| K trial | Pathway | Trial plans | 78/80 | 1.00 (0.94 to 1.00), 60 | 0.83 (0.59 to 0.95), 18 | 0.93 (0.84 to 0.97), 64 | 1.00 (0.76 to 1.00), 14 |
| K trial | Early | No trial plans | 42/42 | 1.00 (0.60 to 1.00), 11 | 0.98 (0.83 to 1.00), 31 | 0.76 (0.41 to 0.94), 14 | 1.00 (0.85 to 1.00), 28 |
| K trial | Early | Trial plans | 58/59 | 0.98 (0.87 to 1.00), 40 | 1.00 (0.79 to 1.00), 18 | 1.00 (0.91 to 1.00), 39 | 0.97 (0.75 to 1.00), 19 |
| K trial | Legacy | No trial plans | 20/20 | 1.00 (0.21 to 1.00), 1 | 0.99 (0.74 to 1.00), 19 | 0.50 (0.09 to 0.91), 2 | 1.00 (0.75 to 1.00), 18 |
| K trial | Legacy | Trial plans | 27/28 | 1.00 (0.80 to 1.00), 16 | 1.00 (0.74 to 1.00), 11 | 1.00 (0.80 to 1.00), 16 | 1.00 (0.74 to 1.00), 11 |
| Any trial | Overall | All | 277/280 | 1.00 (0.98 to 1.00), 193 | 0.95 (0.87 to 0.98), 84 | 0.93 (0.89 to 0.96), 207 | 1.00 (0.94 to 1.00), 70 |
| Any trial | Pathway | No trial plans | 51/51 | 1.00 (0.86 to 1.00), 26 | 0.98 (0.82 to 1.00), 25 | 0.92 (0.75 to 0.98), 29 | 1.00 (0.85 to 1.00), 22 |
| Any trial | Pathway | Trial plans | 79/80 | 1.00 (0.95 to 1.00), 67 | 0.79 (0.50 to 0.93), 12 | 0.94 (0.86 to 0.98), 71 | 1.00 (0.68 to 1.00), 8 |
| Any trial | Early | No trial plans | 41/42 | 1.00 (0.78 to 1.00), 20 | 0.95 (0.76 to 0.99), 21 | 0.80 (0.56 to 0.92), 24 | 1.00 (0.82 to 1.00), 17 |
| Any trial | Early | Trial plans | 59/59 | 1.00 (0.93 to 1.00), 50 | 0.96 (0.64 to 1.00), 9 | 0.98 (0.90 to 1.00), 51 | 1.00 (0.68 to 1.00), 8 |
| Any trial | Legacy | No trial plans | 20/20 | 1.00 (0.71 to 1.00), 10 | 0.97 (0.65 to 1.00), 10 | 0.87 (0.58 to 0.97), 12 | 1.00 (0.68 to 1.00), 8 |
| Any trial | Legacy | Trial plans | 27/28 | 1.00 (0.83 to 1.00), 20 | 1.00 (0.65 to 1.00), 7 | 1.00 (0.83 to 1.00), 20 | 1.00 (0.65 to 1.00), 7 |
| New trial | Overall | All | 277/280 | 0.97 (0.92 to 0.99), 108 | 0.96 (0.91 to 0.99), 169 | 0.89 (0.81 to 0.94), 114 | 0.99 (0.95 to 1.00), 163 |
| New trial | Pathway | No trial plans | 51/51 | 0.95 (0.77 to 0.99), 20 | 0.99 (0.84 to 1.00), 31 | 0.95 (0.75 to 0.99), 19 | 0.99 (0.85 to 1.00), 32 |
| New trial | Pathway | Trial plans | 79/80 | 0.96 (0.83 to 0.99), 31 | 0.95 (0.85 to 0.99), 48 | 0.91 (0.76 to 0.97), 33 | 0.98 (0.89 to 1.00), 46 |
| New trial | Early | No trial plans | 41/42 | 0.96 (0.70 to 1.00), 14 | 0.95 (0.76 to 0.99), 27 | 0.79 (0.52 to 0.92), 14 | 0.99 (0.81 to 1.00), 27 |

| Outcome | Stratum | Intent | Resolved / total | Se (95% CI), n | Sp (95% CI), n | PPV (95% CI), n | NPV (95% CI), n |
|---|---|---|---|---|---|---|---|
| New trial | Early | Trial plans | 58/59 | 1.00 (0.87 to 1.00), 25 | 0.94 (0.79 to 0.99), 33 | 0.89 (0.73 to 0.96), 28 | 1.00 (0.86 to 1.00), 30 |
| New trial | Legacy | No trial plans | 20/20 | 1.00 (0.70 to 1.00), 9 | 0.98 (0.66 to 1.00), 11 | 0.90 (0.60 to 0.98), 10 | 1.00 (0.68 to 1.00), 10 |
| New trial | Legacy | Trial plans | 28/28 | 1.00 (0.70 to 1.00), 9 | 0.93 (0.73 to 0.99), 19 | 0.90 (0.60 to 0.98), 10 | 1.00 (0.82 to 1.00), 18 |
| Trial plans | Overall | All | 97/100 | 0.95 (0.83 to 0.98), 50 | 1.00 (0.91 to 1.00), 47 | 1.00 (0.91 to 1.00), 48 | 0.97 (0.86 to 0.99), 49 |
| Trial plans | Pathway | No trial plans | 17/17 | 0.00 (0.00 to 0.79), 1 | 1.00 (0.81 to 1.00), 16 | Not estimable: zero denominator | 0.94 (0.73 to 0.99), 17 |
| Trial plans | Pathway | Trial plans | 16/17 | 1.00 (0.81 to 1.00), 16 | Not estimable: zero denominator | 1.00 (0.81 to 1.00), 16 | Not estimable: zero denominator |
| Trial plans | Early | No trial plans | 16/17 | Not estimable: zero denominator | 1.00 (0.81 to 1.00), 16 | Not estimable: zero denominator | 1.00 (0.81 to 1.00), 16 |
| Trial plans | Early | Trial plans | 16/17 | 1.00 (0.81 to 1.00), 16 | Not estimable: zero denominator | 1.00 (0.81 to 1.00), 16 | Not estimable: zero denominator |
| Trial plans | Legacy | No trial plans | 16/16 | 0.00 (0.00 to 0.79), 1 | 1.00 (0.80 to 1.00), 15 | Not estimable: zero denominator | 0.94 (0.72 to 0.99), 16 |
| Trial plans | Legacy | Trial plans | 16/16 | 1.00 (0.81 to 1.00), 16 | Not estimable: zero denominator | 1.00 (0.81 to 1.00), 16 | Not estimable: zero denominator |
| New protocol | Overall | All | 60/60 | 0.95 (0.77 to 0.99), 21 | 1.00 (0.91 to 1.00), 39 | 1.00 (0.84 to 1.00), 20 | 0.97 (0.86 to 0.99), 40 |
| Original K | Overall | All | 60/60 | 1.00 (0.91 to 1.00), 39 | 0.95 (0.77 to 0.99), 21 | 0.97 (0.86 to 0.99), 40 | 1.00 (0.84 to 1.00), 20 |
| Developed from K | Overall | All | 60/60 | 0.93 (0.74 to 0.98), 22 | 0.74 (0.57 to 0.85), 38 | 0.69 (0.50 to 0.83), 30 | 0.94 (0.79 to 0.99), 30 |
| Developed from K | Pathway | Trial plans | 30/30 | 1.00 (0.72 to 1.00), 11 | 0.83 (0.59 to 0.94), 19 | 0.81 (0.53 to 0.94), 14 | 1.00 (0.77 to 1.00), 16 |
| Developed from K | Earlier | Trial plans | 30/30 | 0.85 (0.56 to 0.96), 11 | 0.66 (0.44 to 0.83), 19 | 0.59 (0.35 to 0.79), 16 | 0.89 (0.63 to 0.97), 14 |

### Table C. Contemporaneous Required versus Not Allowed comparison

Leadership within five years of K start among all eligible awardees. Risk differences and confidence limits are percentage points. The second panel gives the share of awarded abstracts classified as proposing the awardee's own trial.

**A. Five-year leadership**

| Mechanism | Outcome | Group sizes, n | Required (%) | Not Allowed (%) | RD (95% CI) |
|---|---|---|---|---|---|
| K08 | New-trial leadership | 40/635 | 57.5 | 7.1 | 50.4 (34.9 to 64.5) |
| K08 | Any-trial leadership, including the K trial | 40/635 | 85.0 | 7.6 | 77.4 (63.2 to 85.6) |
| K23 | New-trial leadership | 358/467 | 33.8 | 19.5 | 14.3 (8.2 to 20.4) |
| K23 | Any-trial leadership, including the K trial | 358/467 | 84.1 | 20.8 | 63.3 (57.6 to 68.2) |

Eligibility: Main eligibility. Population: All eligible awardees. Method: Unadjusted. Group sizes: Required / Not Allowed.

**B. Trial plans in awarded abstracts**

| Mechanism | Pathway | Awardees, n | Trial plans, n | Trial plans (%) |
|---|---|---|---|---|
| K08 | Not Allowed | 635 | 6 | 0.9 |
| K08 | Required | 41 | 35 | 85.4 |
| K23 | Not Allowed | 467 | 24 | 5.1 |
| K23 | Required | 358 | 323 | 90.2 |

Instrument estimates and human-corrected estimates use the outcome definitions and validation procedures described above.

**A. Leadership estimates**

| Variant | Quantity | Estimate (95% CI) |
|---|---|---|
| Uncorrected (language model) | K-trial DID | 18.6 (12.0 to 24.9) |
| Uncorrected (language model) | New-trial DID | −2.8 (−11.2 to 6.8) |
| Uncorrected (language model) | K-trial minus new-trial DID | 21.3 (10.9 to 30.7) |
| Uncorrected (language model) | Any-trial DID | 10.8 (2.2 to 19.2) |
| Uncorrected (language model) | Neither-state DID | −10.8 (−19.2 to −2.2) |
| Uncorrected (language model) | K-only-state DID | 13.5 (6.9 to 19.6) |
| Uncorrected (language model) | New-only-state DID | −7.8 (−13.9 to −1.4) |
| Uncorrected (language model) | Both-state DID | 5.0 (−1.0 to 11.2) |
| Primary correction (centered prior) | K-trial DID | 13.7 (5.7 to 21.2) |
| Primary correction (centered prior) | New-trial DID | −3.8 (−14.4 to 6.9) |
| Primary correction (centered prior) | K-trial minus new-trial DID | 17.4 (5.3 to 29.7) |
| Primary correction (centered prior) | Any-trial DID | 4.2 (−5.9 to 14.0) |
| Primary correction (centered prior) | Neither-state DID | −4.2 (−14.0 to 5.9) |
| Primary correction (centered prior) | K-only-state DID | 8.0 (−0.9 to 16.5) |
| Primary correction (centered prior) | New-only-state DID | −9.4 (−16.7 to −2.1) |
| Primary correction (centered prior) | Both-state DID | 5.7 (−1.0 to 12.5) |
| Jeffreys prior | K-trial DID | 15.1 (1.1 to 28.8) |
| Jeffreys prior | New-trial DID | −3.4 (−18.2 to 11.5) |
| Jeffreys prior | K-trial minus new-trial DID | 18.5 (1.2 to 36.1) |
| Jeffreys prior | Any-trial DID | 5.6 (−10.4 to 22.2) |
| Jeffreys prior | New-only-state DID | −9.5 (−20.3 to 1.6) |
| Swapped reference rater | K-trial DID | 13.6 (5.4 to 21.6) |
| Swapped reference rater | New-trial DID | −3.8 (−13.9 to 7.0) |
| Swapped reference rater | K-trial minus new-trial DID | 17.4 (4.5 to 29.0) |
| Swapped reference rater | Any-trial DID | 4.3 (−5.9 to 14.2) |
| Swapped reference rater | New-only-state DID | −9.3 (−16.6 to −1.6) |
| Rater A for double-rated records | K-trial DID | 14.6 (6.4 to 22.8) |
| Rater A for double-rated records | New-trial DID | −3.3 (−12.8 to 7.1) |
| Rater A for double-rated records | K-trial minus new-trial DID | 17.9 (5.7 to 29.3) |
| Rater A for double-rated records | Any-trial DID | 4.2 (−5.5 to 13.8) |
| Rater A for double-rated records | New-only-state DID | −10.3 (−17.5 to −3.1) |
| Rater B for double-rated records | K-trial DID | 13.7 (5.3 to 21.8) |
| Rater B for double-rated records | New-trial DID | −3.7 (−13.8 to 6.6) |
| Rater B for double-rated records | K-trial minus new-trial DID | 17.4 (5.2 to 29.1) |
| Rater B for double-rated records | Any-trial DID | 4.5 (−5.6 to 14.4) |
| Rater B for double-rated records | New-only-state DID | −9.2 (−16.5 to −1.9) |
| Trial plans also corrected, periods pooled | K-trial DID | 11.7 (4.6 to 19.6) |
| Trial plans also corrected, periods pooled | New-trial DID | −1.8 (−11.5 to 7.5) |
| Trial plans also corrected, periods pooled | K-trial minus new-trial DID | 13.5 (2.1 to 25.4) |
| Trial plans also corrected, periods pooled | Any-trial DID | 6.5 (−2.3 to 15.6) |
| Trial plans also corrected, periods pooled | New-only-state DID | −5.2 (−13.2 to 2.5) |
| Trial plans also corrected, by period | K-trial DID | 10.3 (−1.7 to 20.0) |

| Variant | Quantity | Estimate (95% CI) |
|---|---|---|
| Trial plans also corrected, by period | New-trial DID | −4.5 (−14.6 to 5.5) |
| Trial plans also corrected, by period | K-trial minus new-trial DID | 14.8 (0.9 to 27.9) |
| Trial plans also corrected, by period | Any-trial DID | 1.5 (−10.7 to 12.6) |
| Trial plans also corrected, by period | New-only-state DID | −8.8 (−16.1 to −1.4) |
| Correction probabilities pooled across periods | K-trial DID | 16.8 (9.7 to 23.4) |
| Correction probabilities pooled across periods | New-trial DID | −2.4 (−11.0 to 7.2) |
| Correction probabilities pooled across periods | K-trial minus new-trial DID | 19.2 (8.6 to 29.8) |
| Correction probabilities pooled across periods | Any-trial DID | 9.6 (1.5 to 17.9) |
| Correction probabilities pooled across periods | Neither-state DID | −9.6 (−17.9 to −1.5) |
| Correction probabilities pooled across periods | K-only-state DID | 11.9 (5.2 to 18.7) |
| Correction probabilities pooled across periods | New-only-state DID | −7.2 (−13.5 to −0.8) |
| Correction probabilities pooled across periods | Both-state DID | 4.9 (−0.9 to 11.0) |

**B. Scientific continuity**

| Lineage quantity | Estimate (95% CI) |
|---|---|
| Awardee share: post | 17.1 (11.7 to 22.4) |
| Awardee share: early | 17.1 (10.5 to 23.2) |
| Awardee share: legacy | 20.7 (11.1 to 31.7) |
| Awardee difference: post minus early | 0.0 (−8.2 to 8.5) |
| Awardee difference: post minus legacy | −3.6 (−15.2 to 7.2) |
| Pair share: post | 44.2 (29.5 to 56.8) |
| Pair share: early | 38.3 (23.6 to 52.5) |
| Pair share: legacy | 38.4 (19.4 to 58.8) |

### Table E. Composition checks

The first panel reports baseline covariates by trial plans and announcement regime, with the intent-by-period DID for each covariate. Binary covariate means are percentages, and covariate DIDs and confidence limits are percentage points. Institutional size is the log number of cohort K awards. The second panel removes the dedicated-pathway excess intent membership from the top or bottom of the outcome distribution. This trimming scenario assumes added membership without displacement and an unchanged comparator, so it does not bound arbitrary funding selection. Trimming bounds and their confidence limits are percentage points. The third panel uses uncorrected labels: covariate-adjusted and reweighted DIDs, and new-trial DIDs defined by scientific distinctness from the K plan alone (ignoring K-grant links) or by absence of a K-grant link alone.

**A. Covariate balance**

| Covariate | Earlier, intent | Pathway, intent | Earlier, no intent | Pathway, no intent | DID (95% CI) |
|---|---|---|---|---|---|
| Prior non-K trial leadership | 18.1 | 20.2 | 10.7 | 9.8 | 3.0 (−3.5 to 10.8) |
| Prior NIH award as PI | 21.3 | 22.8 | 21.6 | 16.5 | 7.5 (0.1 to 15.2) |
| Prior NIH research grant | 8.5 | 8.1 | 8.2 | 5.1 | 2.9 (−2.2 to 7.8) |
| Log number of cohort K awards at the institution | 3.641 | 3.504 | 3.750 | 3.803 | −0.210 (−0.453 to 0.059) |

Binary covariates: percentages. DID and CI: percentage points. Institutional size: log units. Mechanism: K23.

**B. Excess-share trimming scenario**

| Outcome | Bound | Estimate (95% CI) |
|---|---|---|
| New-trial leadership | Lower | −11.2 (−24.1 to −0.1) |
| New-trial leadership | Upper | 0.0 (−9.5 to 11.0) |
| K-trial leadership within five years | Lower | 14.7 (5.0 to 23.0) |
| K-trial leadership within five years | Upper | 26.0 (16.3 to 37.4) |

Mechanism: K23. Trimmed (%): 10.2.

**C. Uncorrected composition and classification sensitivity DIDs**

| Outcome | Model | Estimate (95% CI) |
|---|---|---|
| New-trial leadership using scientific distinctness alone | Fiscal-year and institute adjustment | −3.1 (−11.9 to 6.3) |
| New-trial leadership using absence of a K-grant link alone | Fiscal-year and institute adjustment | −1.8 (−10.8 to 6.7) |
| New-trial leadership | Adjusted for prior trial, prior NIH awards, institution size | −3.7 (−12.8 to 6.1) |
| K-trial leadership within five years | Adjusted for prior trial, prior NIH awards, institution size | 18.7 (12.1 to 25.1) |
| New-trial leadership | Inverse-probability weighted to earlier covariates, with institute fixed effects | −3.1 (−11.5 to 5.5) |
| K-trial leadership within five years | Inverse-probability weighted to earlier covariates, with institute fixed effects | 18.4 (11.5 to 24.7) |

Analysis: Historical DID. Mechanism: K23. N: 1847.

### Table F. Comparison-group, calendar, and pandemic checks

All outcomes use five years from K start and fiscal-year and institute adjustment. The first panel is human corrected, computed in the same 2,000 institution-bootstrap draws as the primary estimates, so its full-sample rows equal the primary estimates. The comparison restricted to fiscal years 2018–2019 in both regimes aligns calendar time, including pandemic exposure, and consists mostly of fiscal year 2018 legacy and fiscal year 2019 dedicated-pathway awards. It does not separate policies that took effect for applications due from January 25, 2018, which applied to dedicated-pathway but not legacy applications, and it leaves selection into announcements unresolved. The second panel repeats the restricted comparisons with uncorrected (instrument) labels and 1,000 draws. The third panel, also uncorrected, drops trial starts in 2020–2021 while retaining all awardees. Because dedicated-pathway awardees started later, this exclusion removes more of their follow-up window than of earlier awardees' windows and is biased toward smaller estimates.

**A. Human-corrected comparisons**

| Comparison | Outcome | DID (95% CI) | Awardees, n |
|---|---|---|---|
| Full sample | K-trial DID | 13.7 (5.7 to 21.2) | 1847 |
| Full sample | New-trial DID | −3.8 (−14.4 to 6.9) | 1847 |
| Full sample | K-trial minus new-trial DID | 17.4 (5.3 to 29.7) | 1847 |
| Legacy awards as comparator | K-trial DID | 13.5 (−0.3 to 27.3) | 1055 |
| Legacy awards as comparator | New-trial DID | −4.6 (−19.2 to 10.8) | 1055 |
| Legacy awards as comparator | K-trial minus new-trial DID | 18.1 (−1.0 to 34.6) | 1055 |
| Fiscal years 2018-2019 in both regimes | K-trial DID | 12.9 (−3.9 to 29.1) | 494 |
| Fiscal years 2018-2019 in both regimes | New-trial DID | −7.6 (−25.6 to 9.6) | 494 |
| Fiscal years 2018-2019 in both regimes | K-trial minus new-trial DID | 20.5 (−0.5 to 41.8) | 494 |

**B. Uncorrected restricted comparisons**

| Comparison | Outcome | DID (95% CI) | Earlier / pathway, n |
|---|---|---|---|
| Fiscal year 2018–2019 legacy versus all pathway awards | K trial | 18.2 (3.6 to 31.7) | 218 / 837 |
| Fiscal year 2018–2019 legacy versus all pathway awards | New trial | −6.1 (−19.6 to 5.8) | 218 / 837 |
| Fiscal year 2018–2019 in both regimes | K trial | 17.9 (1.8 to 34.6) | 218 / 276 |
| Fiscal year 2018–2019 in both regimes | New trial | −6.5 (−25.3 to 8.2) | 218 / 276 |

Mechanism: K23.

**C. Excluding trial starts in 2020-2021 (uncorrected)**

| Outcome | DID (95% CI) |
|---|---|
| K trial | 7.7 (0.6 to 14.4) |
| New trial | 0.7 (−6.8 to 7.9) |
| K minus new | 7.0 (−1.9 to 16.9) |

Awardees, n: 1847.

### Table G. Cohort trends and bounded drift

Human corrected, in the primary 2,000 institution-bootstrap draws. Fiscal-year 2014–2017 intent-specific slopes adjust for year and institute. Subtracting this slope before refitting the DID gives the extrapolated estimate. The drift envelope permits additional intent-specific annual drift after fiscal year 2017, shifting each draw by its own drift exposure. Breakdown denotes the annual bound where the interval first includes zero. These additive-bias envelopes are not formal robust event-study confidence sets. Early-cohort follow-up extends into the policy era, precluding uncontaminated earlier outcome leads. The placebo compares the trial-plan gap in fiscal years 2016–2017 with 2014–2015 among early awards only. The mean drift exposure is 2.76 years, so the breakdown approximately equals the lower confidence limit divided by this exposure.

##### A. Cohort trends

| Model | Outcome | Estimate (95% CI) | Units |
|---|---|---|---|
| Extrapolated early-cohort trend | K-trial DID | 4.1 (−17.7 to 26.8) | Percentage points |
| Extrapolated early-cohort trend | New-trial DID | −16.8 (−39.3 to 6.1) | Percentage points |
| Extrapolated early-cohort trend | K-trial minus new-trial DID | 21.0 (−11.5 to 55.4) | Percentage points |
| Differential early-cohort slope (per year) | K-trial DID | 2.4 (−3.0 to 7.9) | Points per year |
| Differential early-cohort slope (per year) | New-trial DID | 3.3 (−1.8 to 8.3) | Points per year |
| Differential early-cohort slope (per year) | K-trial minus new-trial DID | −0.9 (−8.7 to 6.3) | Points per year |
| Early-cohort placebo, fiscal years 2016–2017 versus 2014–2015 (uncorrected) | K-trial DID | 7.7 (−1.7 to 19.2) | Percentage points |

##### B. Bounded-drift envelopes

| Outcome | Annual drift bound | DID envelope (95%) | Interval breakdown, points/year |
|---|---|---|---|
| K-trial DID | 0 | 13.7 (5.7 to 21.2) | 2.03 |
| K-trial DID | 0.5 | 13.7 (4.3 to 22.6) | 2.03 |
| K-trial DID | 1 | 13.7 (2.9 to 24.0) | 2.03 |
| K-trial DID | 2 | 13.7 (0.1 to 26.7) | 2.03 |
| New-trial DID | 0 | −3.8 (−14.4 to 6.9) | 0 |
| New-trial DID | 0.5 | −3.8 (−15.7 to 8.3) | 0 |
| New-trial DID | 1 | −3.8 (−17.1 to 9.7) | 0 |
| New-trial DID | 2 | −3.8 (−19.9 to 12.4) | 0 |
| K-trial minus new-trial DID | 0 | 17.4 (5.3 to 29.7) | 1.95 |
| K-trial minus new-trial DID | 0.5 | 17.4 (4.0 to 31.1) | 1.95 |
| K-trial minus new-trial DID | 1 | 17.4 (2.6 to 32.5) | 1.95 |
| K-trial minus new-trial DID | 2 | 17.4 (−0.1 to 35.2) | 1.95 |

### Table H. Grant-link decomposition and PI of record

Historical K-grant matching uses the identification module, with no free-text-only qualification. No recorded serial means no identification-module link, rather than a search proving its absence throughout free text. Component awardee shares can overlap across trials, so component DIDs need not sum to the total. PI diagnostics use current fields on all linked candidate records, including observational records and records outside the outcome window. Awardee PI takes precedence when co-listed with another person. Record categories are exclusive, while awardee categories may overlap. Shares use all intent awardees as denominators and do not establish initial PI designation or actual responsibility. Component contrasts use fiscal-year and institute adjustment with 1,000 institution draws.

**A. Grant-link components**

| Stratum | Trial plans | Component | Awardees | Events | Share (%) |
|---|---|---|---|---|---|
| Early | No | Identification module | 491 | 15 | 3.1 |
| Early | No | Free text only | 491 | 0 | 0.0 |
| Early | No | Judged from plan, no recorded serial | 491 | 9 | 1.8 |
| Early | Yes | Identification module | 301 | 122 | 40.5 |
| Early | Yes | Free text only | 301 | 0 | 0.0 |
| Early | Yes | Judged from plan, no recorded serial | 301 | 50 | 16.6 |
| Legacy | No | Identification module | 144 | 2 | 1.4 |
| Legacy | No | Free text only | 144 | 0 | 0.0 |
| Legacy | No | Judged from plan, no recorded serial | 144 | 1 | 0.7 |
| Legacy | Yes | Identification module | 74 | 33 | 44.6 |
| Legacy | Yes | Free text only | 74 | 0 | 0.0 |
| Legacy | Yes | Judged from plan, no recorded serial | 74 | 11 | 14.9 |
| Pathway | No | Identification module | 491 | 16 | 3.3 |
| Pathway | No | Free text only | 491 | 0 | 0.0 |
| Pathway | No | Judged from plan, no recorded serial | 491 | 10 | 2.0 |
| Pathway | Yes | Identification module | 346 | 205 | 59.2 |
| Pathway | Yes | Free text only | 346 | 0 | 0.0 |
| Pathway | Yes | Judged from plan, no recorded serial | 346 | 60 | 17.3 |

**B. Adjusted component contrasts**

| Outcome | DID (95% CI) |
|---|---|
| K trial with identification-module link | 17.8 (10.8 to 24.8) |
| K trial with free-text-only link | 0.0 (0.0 to 0.0) |
| K trial judged from plan without serial | 0.8 (−4.2 to 6.0) |

N: 1847.

**C. Current PI of record**

| Stratum | PI of record | Intent awardees | Any linked record | Category awardees | Share (%) | Candidate records |
|---|---|---|---|---|---|---|
| Early | Awardee | 301 | 141 | 134 | 44.5 | 149 |
| Early | Another person | 301 | 141 | 1 | 0.3 | 1 |
| Early | No PI named | 301 | 141 | 7 | 2.3 | 7 |
| Legacy | Awardee | 74 | 42 | 38 | 51.4 | 40 |
| Legacy | Another person | 74 | 42 | 2 | 2.7 | 2 |
| Legacy | No PI named | 74 | 42 | 2 | 2.7 | 2 |
| Pathway | Awardee | 346 | 236 | 221 | 63.9 | 248 |
| Pathway | Another person | 346 | 236 | 4 | 1.2 | 4 |
| Pathway | No PI named | 346 | 236 | 16 | 4.6 | 16 |

### Table I. Leadership by period and trial plans

Instrument-measured leadership within five years of K start. Percentages count awardees. K trials either cite the K grant or are judged to match the K plan. The second panel describes the first K trial starting within five years of K start for each trial-plan awardee with such a trial, by period. Results reporting is in Table J.

**A. Leadership by period**

| Period | Trial plans | Awardees, n | K trial (%) | New trial (%) | Any trial (%) |
|---|---|---|---|---|---|
| Earlier | Trial plans | 375 | 54.4 | 37.6 | 69.9 |
| Earlier | No trial plans | 635 | 4.1 | 19.1 | 20.9 |
| Pathway | Trial plans | 346 | 73.1 | 33.5 | 81.5 |
| Pathway | No trial plans | 491 | 5.1 | 18.7 | 23.0 |

Mechanism: K23.

**B. First K-trial characteristics**

| Characteristic | Earlier | Pathway |
|---|---|---|
| First K trials, n | 204 | 253 |
| Median enrollment | 60 | 58 |
| Enrollment ≥100 (%) | 18.1 | 24.1 |
| Randomized (%) | 77.9 | 68.0 |
| At least three sites (%) | 3.9 | 5.5 |
| Phase 2 or higher (%) | 16.2 | 13.8 |
| Completed (%) | 84.3 | 66.0 |
| Terminated or withdrawn (%) | 9.8 | 6.3 |
| Ongoing (%) | 1.5 | 22.5 |
| Registered within 12 months of start (%) | 98.0 | 99.2 |

Mechanism: K23.

### Table J. Trial-level reporting

The reported outcome is trial-level first results submission within twelve months of primary completion. Due trials were COMPLETED or TERMINATED with positive ACTUAL enrollment, with primary completion at least twelve calendar months before September 27, 2026. Month-only dates use the first day. Primary-completion ACTUAL/ESTIMATED type was not retrieved and was not imposed. UNKNOWN trials with past completion dates are reported separately and excluded from due denominators. Each eligible trial appears once, with no duplicates across awardees or strata. Both trial metrics use first results submission, not public posting after quality control. Ever submission includes reports after twelve months and remains subject to unequal follow-up. All intervals use institution resampling.

##### A. Trial-level reporting denominators and shares

| Characteristic | Early | Legacy | Pathway |
|---|---|---|---|
| Unique K trials | 183 | 46 | 289 |
| Due trials | 165 | 40 | 170 |
| Terminated due trials | 13 | 2 | 10 |
| Submitted within 12 months, n | 42 | 18 | 79 |
| Submitted within 12 months (%) | 25.5 | 45.0 | 46.5 |
| Ever submitted, n | 93 | 35 | 136 |
| Ever submitted (%) | 56.4 | 87.5 | 80.0 |
| UNKNOWN with past primary completion | 8 | 2 | 11 |
| Missing primary-completion date | 0 | 0 | 0 |

##### B. Trial-level contrasts

| Outcome | Comparison | Difference (95% CI) | Trials, n |
|---|---|---|---|
| Within 12 months | Post minus early | 21.0 (11.3 to 30.9) | 335 |
| Ever submitted | Post minus early | 23.6 (14.1 to 33.9) | 335 |
| Within 12 months | Post minus legacy | 1.5 (−17.1 to 18.7) | 210 |
| Ever submitted | Post minus legacy | −7.5 (−18.7 to 6.0) | 210 |

### Table K. Registration by the follow-up horizon

Both horizons use the same complete-seven-year K23 sample. Symmetric registration requires first submission no later than K start plus the respective horizon. Five-year outcomes are nested within seven-year outcomes, with paired institution draws and fiscal-year and institute adjustment. The rule excludes retrospective registrations after each horizon but cannot recover never-registered or unretrieved trials. Human leadership validation covers five years only.

**A. Registration by horizon**

| Horizon | New-trial DID (95% CI) |
|---|---|
| Five years | −6.3 (−18.2 to 5.2) |
| Seven years | −3.4 (−16.7 to 8.9) |

Mechanism: K23. Registration rule: First submitted by horizon. Earlier / pathway, n: 1010 / 264.

### Table L. Further historical estimates

Historical DIDs adjust for fiscal year and institute. Estimates and confidence limits are percentage points. The drug, biologic or device restriction counts only K trials testing those interventions. Late-start new trials begin at least three years after K start. R01-equivalent awards within five years follow the NIH Data Book definition: DP1, DP2, DP5, R01, R37, R56, RF1, RL1 and U01, plus R35 from NIGMS or NHGRI only. K08 estimates rest on 14 earlier trial-plan awardees.

**A. Estimates**

| Mechanism | Outcome | Model | Estimate (95% CI) | N |
|---|---|---|---|---|
| K23 | K-trial leadership in drug, biologic or device trials | Drug/biologic/device trials only | 2.8 (−2.9 to 8.1) | 1847 |
| K23 | New-trial leadership starting at least three years after K start | Fiscal-year and institute adjustment | −1.4 (−8.4 to 6.8) | 1847 |
| K23 | R01-equivalent award within five years (NIH definition) | Fiscal-year and institute adjustment | −0.7 (−8.1 to 6.7) | 1847 |
| K08 | New-trial leadership | Fiscal-year and institute adjustment | 18.8 (−4.0 to 40.0) | 1515 |
| K08 | K-trial leadership within five years | Fiscal-year and institute adjustment | 2.1 (−29.5 to 45.7) | 1515 |

Analysis: Historical DID.

### Fig A. Registration categories, initial leadership, and recorded initiation among K23 awardees with trial plans

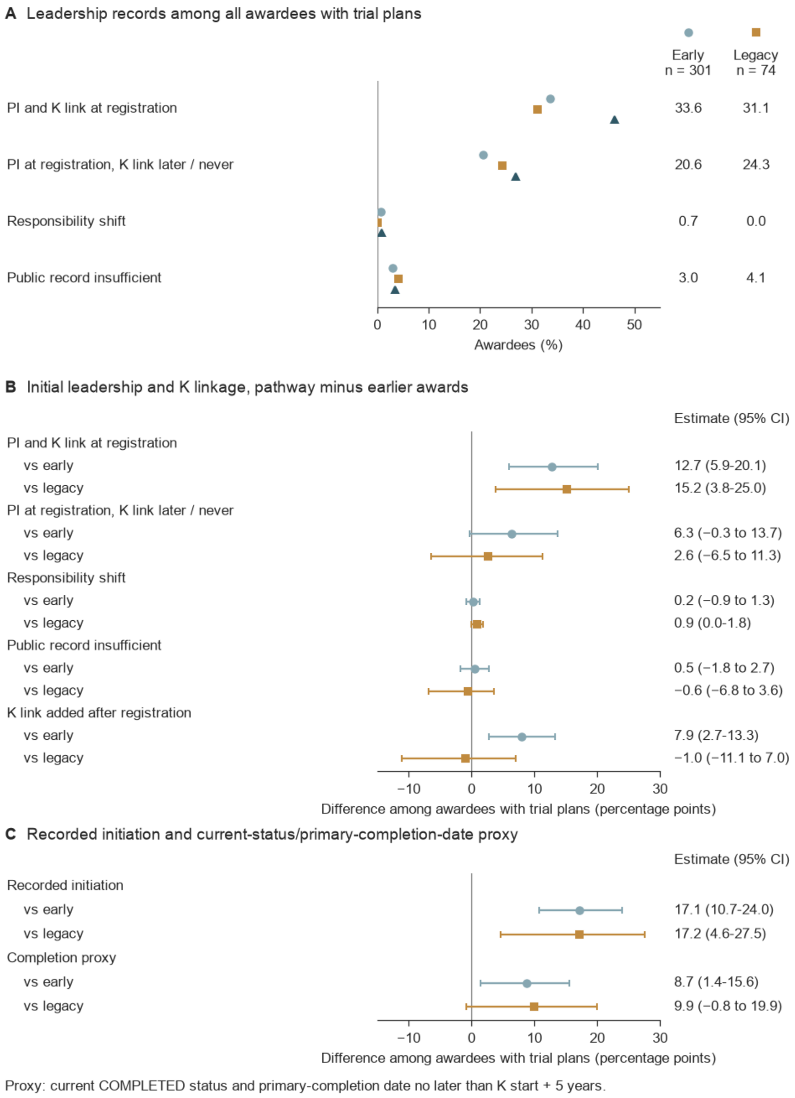


A, Registration categories are exclusive for individual trials, although awardees can contribute trials to several categories. B, Initial leadership and K linkage differences between dedicated-pathway awards and early or legacy awards. C, Recorded initiation and a current-status/primary-completion-date proxy, with all trial-plan awardees as denominators. Recorded initiation means that a K trial was recruiting, enrolling by invitation, active, or completed in any registry version, or reported actual enrollment above zero. It was documented for 52.8%, 52.7%, and 69.9% of early, legacy, and pathway awardees with trial plans, and the completion proxy for 30.9%, 29.7%, and 39.6%. Completion requires current COMPLETED status and primary completion within 5 years of K start, without verifying overall completion. Whiskers show 95% CIs from bootstrap draws resampling institutions. Shares and differences are uncorrected and compare awardees with trial plans across strata. They are not difference-in-differences estimates.

### Fig B. Trial-level results reporting by award stratum

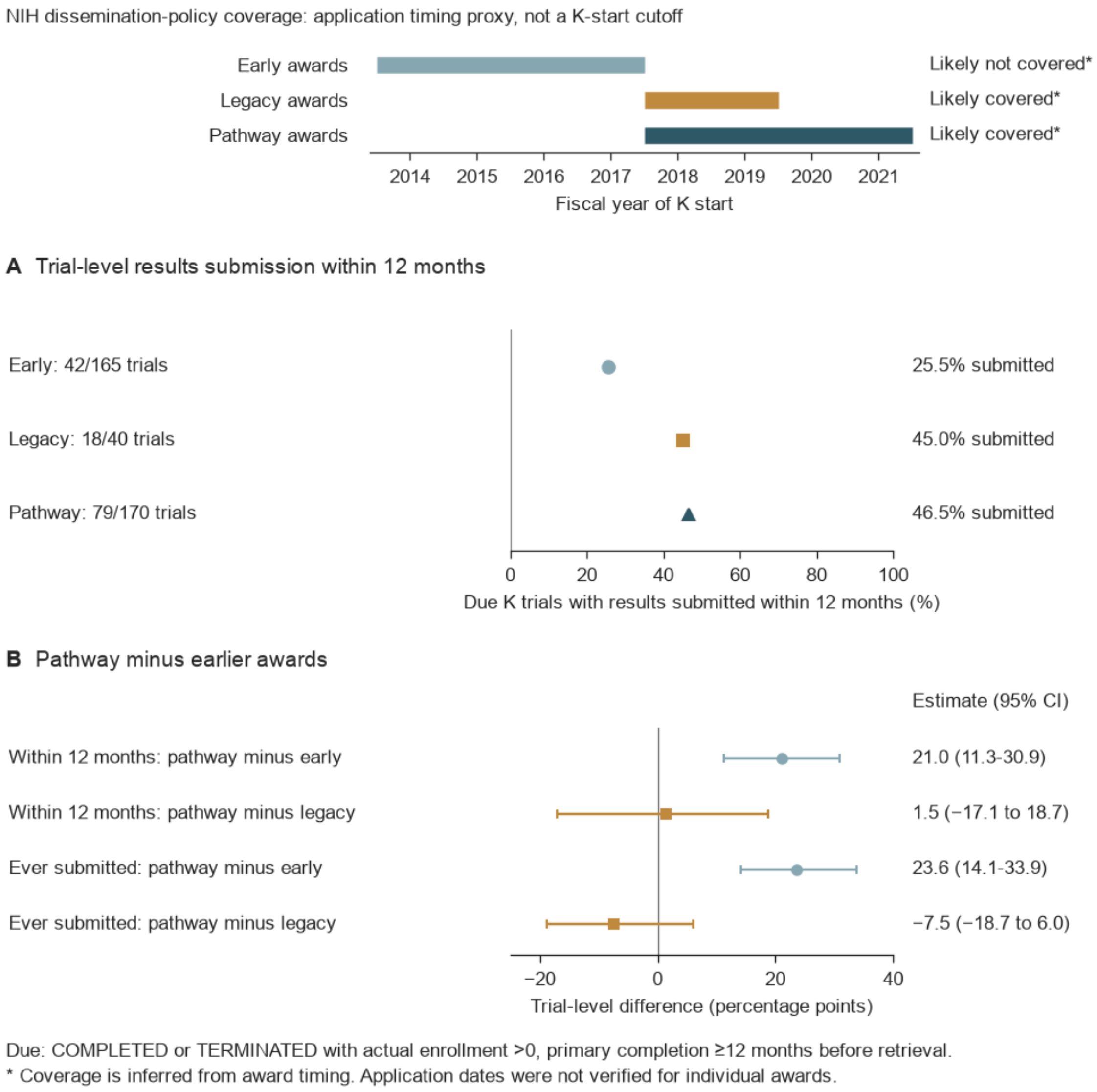


The timeline shows fiscal years of K start. Coverage labels summarize a timing proxy for the NIH dissemination policy, and individual application dates were not verified. A, Results submission within 12 months of primary completion among due trials, which were COMPLETED or TERMINATED with positive actual enrollment and primary completion at least 12 months before retrieval. B, Differences between dedicated-pathway and earlier strata in within-12-month and ever-submitted results. Whiskers show 95% CIs from bootstrap draws resampling institutions.
